# LongcallD: joint calling and phasing of small, structural and mosaic variants from long reads

**DOI:** 10.64898/2026.03.20.713111

**Authors:** Yan Gao, Wen-Wei Liao, Qian Qin, Ira M. Hall, Heng Li

## Abstract

Long-read sequencing is a powerful technique capturing multiple variants within single continuous reads. This length allows individual reads to bridge small and structural variants while carrying crucial phasing information. However, current computational tools treat small variant calling, structural variant (SV) detection and phasing as largely disconnected problems, failing to unleash the full potential of long reads. Here, we present longcallD, a unified framework utilizing local multiple-sequence alignment to simultaneously call and phase small and structural variants. By integrating germline phasing and retrotransposition hallmarks, longcallD also identifies low-fraction mosaic variants and detects mobile element insertions supported by a single read. Compared to existing methods, our unified approach substantially improves SV discovery and mosaic variants accuracy while maintaining competitive small variant calling. We anticipate that longcallD will provide a robust foundation for resolving complex genetic architectures in clinical and evolutionary applications.

## Introduction

Long-read DNA sequencing technologies from Pacific Biosciences (PacBio) and Oxford Nanopore Technologies (ONT) have fundamentally expanded the scope of genomic variant detection^1^. Long reads spanning tens of thousands of base pairs enable direct interrogation of complex genomic regions that remain inaccessible to short-read approaches^2^. The advent of PacBio high-fidelity (HiFi) sequencing and ONT R10.4.1 chemistry has further improved base-level read accuracy, substantially reducing sequencing errors and enhancing variant calling performance^3,4^. Despite these progresses, accurately calling the full spectrum of genetic variation that encompasses single-nucleotide polymorphisms (SNPs), small insertions and deletions (indels), and large structural variants (SVs) from long-read data remains a substantial computational challenge.

With short reads, small variant calling, SV calling and phasing are three independent problems as a standard short read rarely bridges ≥2 variants or spans an entire SV^5^. These problems are thus solved by distinct algorithms. In contrast, a single long read often connects multiple small and structural variants and naturally encodes their phasing. Long reads have unified small/structural variant calling and phasing but this advantage has not been fully utilized. On small variant calling, modern long-read callers including longshot^6^, Clair3^7^ and DeepVariant^8,9^ either natively implement a phasing algorithm or use a third-party phasing tool. However, unaware of SVs, they are sensitive to inconsistent alignment around SVs, especially in low-complexity regions (LCRs), which may lead to spurious variant calls^2,10^. Clair3 and DeepVariant rely on deep learning to filter such errors but without multiple-sequence realignment, they are unable to recover small variants tangled with SVs and thus lose power.

On SV calling, early tools such that DELLY^11^, pbsv, DeBreak^12^, SVIM^13^ and cuteSV^14^ ignore read phasing. SVDSS^15^ and sawfish^16^ apply local realignment or reassembly to phase close SVs but do not leverage phasing in entire reads. Sniffles2^17^ and Severus^18^ optionally use full read phasing derived from third-party tools but without realigning or reassembling reads, they often fail to resolve SVs in LCRs; false SNP calls caused by SVs in turn reduce phasing accuracy and subsequently SV accuracy as well. Sniffles2 is the only SV caller that is developed with mosaic SV calling in mind, but it produced hundreds of false mosaic SV calls as we will show later. In summary, although modern variant callers use phasing to some extent, none of them tightly integrates small variant calling, SV calling and phasing.

To address these challenges, we developed longcallD, a unified framework for the joint calling and phasing of small variants and SVs from long genomic reads. LongcallD explicitly distinguishes clean and noisy genomic regions, applies haplotype-aware multiple sequence alignment (MSA) within noisy regions to derive consensus sequences, and integrates clean- and noisy-region variant calls through an iterative phasing procedure. The phasing information is then leveraged to call large indels and mosaic single-nucleotide variants (SNVs) of low variant allele fraction (VAF). We benchmarked longcallD against state-of-the-art variant callers on HiFi and ONT data from the well-characterized HG002 reference sample^10^ and demonstrated its improved germline variant calling performance especially in tandem repeat regions. We also illustrated longcallD’s utility for accurate mosaic variant detection using tumor–normal mixture experiments.

## Results

### Overview of longcallD

Given a reference genome and a long-read alignment, longcallD produces phased variant calls and can optionally output phased long reads with refined alignments. The overall workflow is illustrated in **Fig. 1**, with algorithmic details described in Methods.

**Fig. 1.**
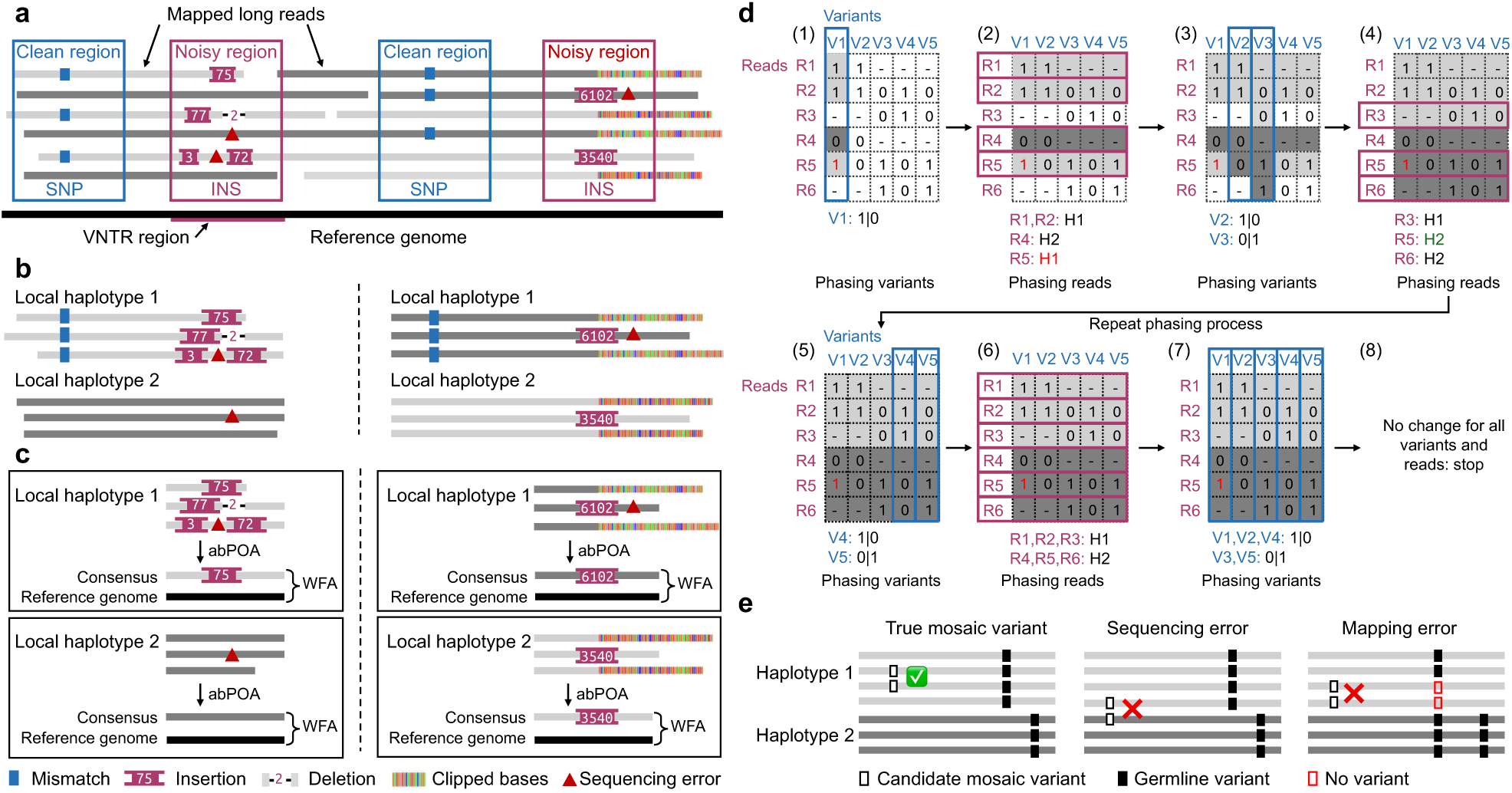
Illustration of the longcallD variant calling workflow. **a**, Classification of candidate regions into clean and noisy categories. **b**, Local haplotagging of long reads using heterozygous variants identified in clean regions. **c**, Haplotype-aware consensus generation and variant calling within noisy regions. **d**, Iterative haplotype extension by alternately traversing variants (blue box) and supporting reads (pink box). The read–variant compatibility matrix indicates whether a read supports the reference allele (0), the variant allele (1), or does not cover the site (–) for a given variant. The red number indicates a sequencing error that initially led to incorrect haplotype assignment but was corrected in a subsequent iteration. (1) The heterozygous variant with the highest coverage (V1) is selected to initialize phasing. (2) Reads covering V1 are assigned to haplotypes according to the allele they support. (3) Variants V2 and V3 are phased based on haplotype assignments of overlapping reads; no read haplotypes change after phasing V2. (4) Following phasing of V3, reads R3 and R6 are assigned haplotypes and the haplotype of R5 is updated. (5) Variants V4 and V5 are phased. (6–7) Read and variant are iteratively traversed for haplotype updating. (8) Iteration terminates when haplotype assignments converge with no further changes. **e**, Detection of low-VAF mosaic variants leveraging haplotype information from long reads. Left, true mosaic variant with supporting reads confined to a single haplotype; middle, false mosaic variant with supporting reads distributed across both haplotypes; right, false mosaic variant with supporting reads inconsistent with phased germline variants.

LongcallD is motivated by the widespread presence of heterozygous SNPs and small indels in human genomes^5^ and by the observation that variant calling errors in long reads are highly enriched in specific genomic contexts. LongcallD identifies noisy regions that comprise (i) homopolymers and short tandem repeats (STRs), where sequencing errors occur more frequently; (ii) Variable number tandem repeats (VNTR), where read alignments are highly inconsistent^19^; and (iii) large insertion or deletion SV regions, where alignments frequently exhibit clipping near SV boundaries. The remaining regions are designated as clean, in which read alignments are generally accurate and concordant. High-confidence germline SNPs and small indels are called directly from clean regions by allele counting.

LongcallD then exploits heterozygous variants in clean regions (allele fraction 0.2–0.8) to locally partition long reads into haplotypes. For each noisy region, haplotype-specific read subsequences are extracted and subjected to MSA to derive haplotype-resolved consensus sequences. In regions lacking nearby heterozygous variants, longcallD performs haplotype-agnostic MSA followed by sequence clustering to distinguish heterozygous from homozygous regions. Consensus sequences are realigned to the reference to generate variant calls, which are subsequently integrated with variants in clean regions through an iterative haplotype-extension procedure to produce jointly phased calls across the genome.

Beyond germline variants, longcallD also calls mosaic variants of low VAF. By leveraging established haplotype information together with a stringent, context-aware filtering strategy, longcallD distinguishes true mosaic mutations from long-read sequencing artifacts. The current implementation focuses on mosaic SNVs and large indels, including mobile element insertions (MEIs) which typically have explicit sequence features like target site duplications (TSDs) and poly(A) tails^20^.

### Benchmark datasets and evaluation framework for germline variants

We evaluated germline small- and structural-variant calling performance using the Genome-in-a-Bottle (GIAB) whole-genome assembly-based benchmark constructed from the T2T HG002 Q100 v1.1 diploid assembly^10^. The GRCh38 version of this benchmark contains 4.6 million small variants and 28,246 large indels. Compared to the previous HG002 SV benchmark v0.6^21^, T2T Q100 v1.1 includes roughly twice as many SVs, with the majority of newly added variants located in LCRs, which are known to be challenging for variant calling^2,10^.

We used one PacBio HiFi (63x) and one ONT R10.4.1 (50x) dataset of the HG002 sample for evaluation. For each dataset, we generated de novo assemblies using long reads only with hifiasm^22,23^ and additionally included the T2T HG002 assembly for comparison. Long reads and assemblies were aligned to GRCh38, and variants were called from both read alignments and assembly alignments. LongcallD was compared with two long-read small variant callers (Clair3^7^ and DeepVariant^9^), eight long-read SV callers (cuteSV^14^, DeBreak^12^, DELLY^11^, pbsv^24^, sawfish^16^, Sniffles2^17^, SVDSS^15^, and SVIM^13^), and four assembly-based variant callers (cuteSV-asm, i.e., cuteSV in assembly mode^14^, dipcall^25^, PAV^26^, and SVIM-asm^13^). Among these, longcallD, dipcall, and PAV report both small variants and SVs.

We measure the small-variant accuracy using hap.py^27^, and the SV accuracy using Truvari^28^. However, upon examining the Truvari reports, we found that Truvari does not require exact allele matches, leading to a substantial underestimation of SV errors. To better characterize SV calling performance in LCRs, we additionally used aardvark^29^, a haplotype-centric benchmarking tool that directly compares haplotype sequences rather than variant representations. Aardvark reports both genotype-level scores and basepair-level scores, the latter allowing partial credit for sequence-level concordance which is an essential property for benchmarking in LCRs.

### Haplotype tagging of noisy regions via flanking heterozygous variants

We first evaluated the effectiveness of noisy-region classification and local haplotype partitioning of longcallD. Only noisy regions overlapping at least one benchmark variant were considered. For HiFi data, 93.4% (673,943 of 721,633) of noisy regions were flanked by at least one heterozygous clean-region variant, enabling haplotype-aware separation of long reads (**Fig. 2a**). For ONT data, this fraction increased to 96.3% (738,826 of 766,898) due to the longer ONT reads which span informative heterozygous sites over larger distances.

**Fig. 2.**
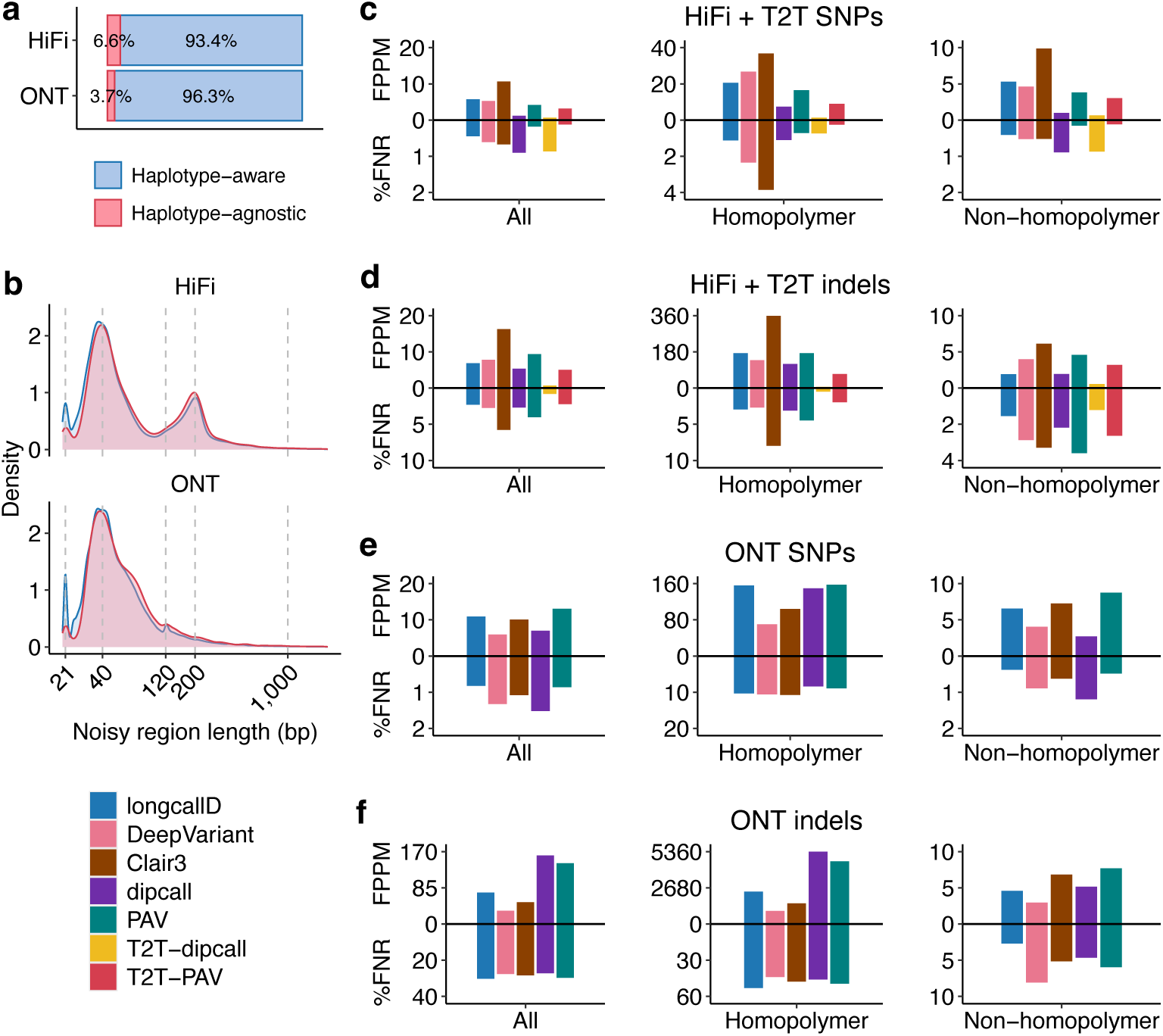
Characterization of noisy regions and small-variant calling accuracy. **a**, Fractions of haplotype-aware and haplotype-agnostic noisy regions identified by longcallD on HG002. **b**, Length distribution of noisy regions identified by longcallD on HG002. **c**, False positives per million bases (FPPM) and false-negative rates (FNR) for SNPs on HG002 HiFi data and corresponding T2T assemblies, evaluated using hap.py. Variants are stratified into all regions, homopolymer regions, and non-homopolymer regions. LongcallD, DeepVariant, and Clair3 were applied to long-read alignments, dipcall and PAV were applied to alignments of long-read assemblies, T2T-dipcall and T2T-PAV were applied to alignments of T2T assemblies. **d**, FPPM and FNR for indels on HG002 HiFi data. **e**, FPPM and FNR for SNPs on HG002 ONT data. **f**, FPPM and FNR for indels on HG002 ONT data.

We next examined the length distribution of noisy regions (**Fig. 2b**). Haplotype-aware and haplotype-agnostic noisy regions exhibited similar length distributions for both technologies, with a pronounced peak at approximately 40 bp. This peak corresponds primarily to homopolymer-associated regions consisting of ∼20-bp homopolymers flanked by two additional 10-bp windows. Although long noisy regions constitute a minority, they are strongly enriched for tandem repeats: 53.9% of HiFi noisy regions ≥200 bp and 58.6% of ONT noisy regions ≥120 bp overlap annotated tandem repeats. These results demonstrate that longcallD effectively leverages long-range haplotype information to resolve most challenging regions encountered in long-read data.

### Comparable small-variant calling accuracy to state-of-the-art methods

We extracted SNPs and small indels (<50 bp) from variant calls generated by longcallD, PAV, and dipcall, and evaluated them together with DeepVariant and Clair3 against the ground-truth call set. Accuracy was assessed using precision, recall, F1 score, false-negative rate (FNR), and false positives per million bases (FPPM), with results stratified into homopolymer and non-homopolymer regions based on GIAB genome stratification v3.6^30^.

Using HiFi reads, all callers exhibited higher error rates in homopolymer regions than in non-homopolymer regions (**Fig. 2c,d**). Among alignment-based callers, longcallD achieved the lowest FNRs and highest F1 scores for both SNPs and indels across all confident regions (**Supplementary Fig. 1a,b**). LongcallD produced fewer false positives and false negatives for SNPs in homopolymer regions, potentially due to MSA, whereas DeepVariant achieved higher accuracy for homopolymer indels probably because it models the homopolymer error profile better with deep learning.

Assembly-based callers dipcall and PAV showed comparable SNP calling accuracy between HiFi and T2T assemblies; however, hifiasm HiFi assembly carried noticeably more indel errors especially in homopolymer regions, while the T2T assembly did not have this issue. Relative to dipcall and PAV applied to the HiFi assembly, longcallD exhibited a lower FNR but a higher FPPM. Overall, with HiFi long reads, longcallD demonstrated small variant calling performance comparable to both alignment-based and assembly-based state-of-the-art methods.

With ONT long reads, all callers called substantially more homopolymer SNP and indel errors compared to HiFi data (**Fig. 2c–f**). LongcallD achieved the lowest FNRs for both SNPs and indels in non-homopolymer regions, whereas DeepVariant and Clair3 showed superior performance in homopolymer regions with better error modeling with deep learning. Using the latest ONT-only assembly generated by hifiasm (ONT)^23^, both dipcall and PAV exhibited higher error rates than when applied to the HiFi assembly, particularly for indels in homopolymer regions (**Supplementary Fig. 1c,d**).

Downsampling experiments demonstrated that longcallD’s small-variant accuracy improved with increasing sequencing depth, with the largest gains between 15× and 20× coverage and saturation beyond ∼30×, particularly in homopolymer regions (**Supplementary Fig. 2**). Similar trends were observed for DeepVariant and Clair3.

### Improved SV calling accuracy in low-complexity regions

For SV calling evaluation, we extracted large indels (length ≥ 30 bp) from variant calls generated by longcallD, PAV, and dipcall, and evaluated them together with other SV-only callers against the ground-truth call set. Accuracy was measured by FNR and false discovery rate (FDR), with SVs stratified into tandem-repeat and non-tandem-repeat (“other”) regions using the GIAB v3.6 annotations^30^.

Although tandem repeat regions comprise only 42 Mb of the total 2.5 Gb confident region, they account for most SV calling errors across all methods (**Fig. 3a,b**). On both HiFi and ONT datasets, longcallD achieved the lowest FDR and FNR among all alignment-based SV callers. For SVs calling in tandem-repeat regions using HiFi data, longcallD attained the best FDR of 3.2% and the best FNR of 9.8% simultaneously, a leap from the second-best FDR of 5.1% (SVDSS) and FNR of 12.3% (sawfish). This demonstrates the power of unified variant calling and phasing.

**Fig. 3.**
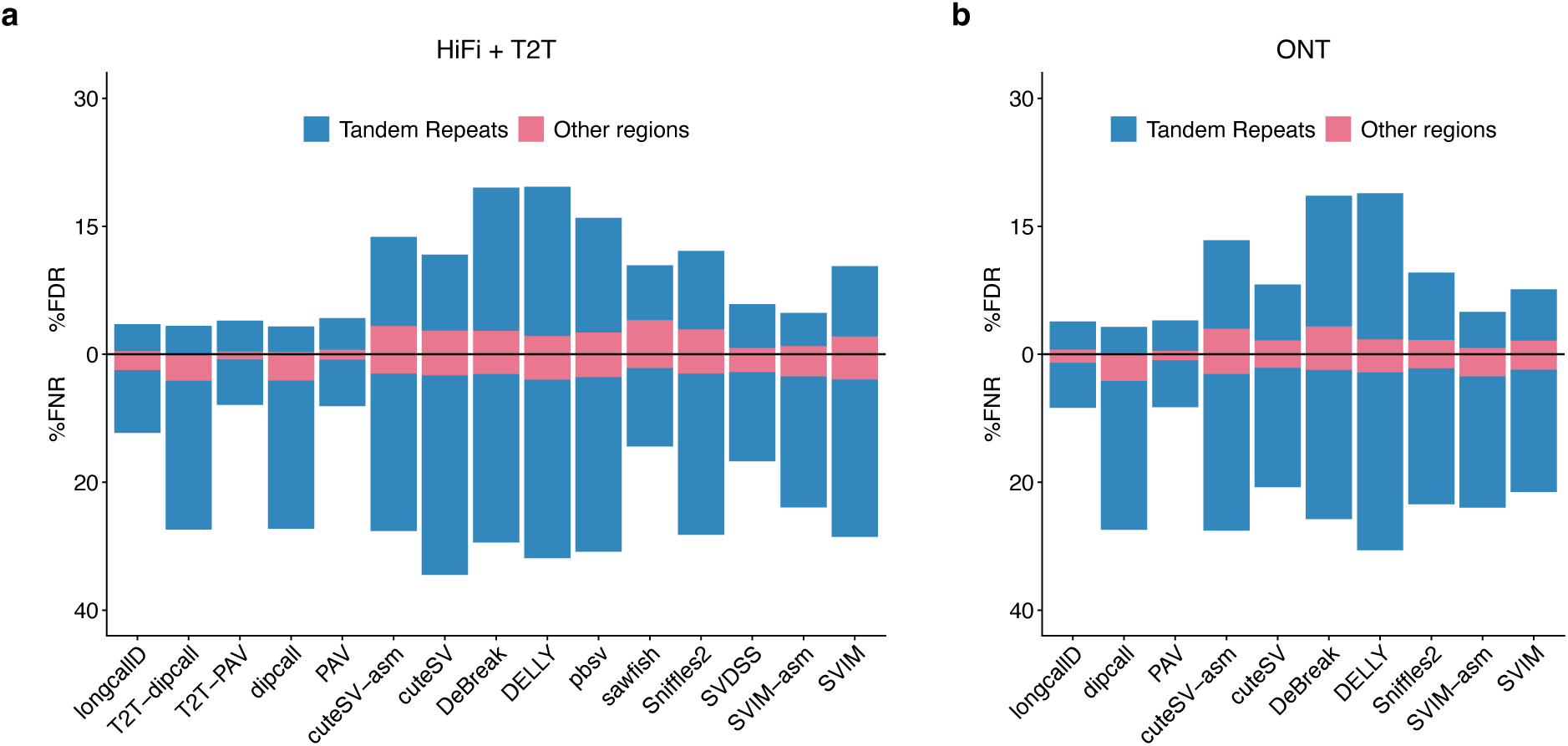
Accuracy of SV calling. **a**, FDR and FNR of SVs on HiFi and T2T data. Metrics were calculated using truvari in “refine” mode, with SVs stratified by genomic context (tandem repeats vs. other regions). Dipcall, PAV, cuteSV-asm, and SVIM-asm were applied to long-read assembly alignments, T2T-dipcall and T2T-PAV were applied to T2T assembly alignments. All other callers were applied to long-read alignments. **b**, FDR and FNR of SVs on ONT data.

Assembly-based callers generally outperformed alignment-based callers. Among them, PAV achieved lower FNRs than dipcall across all assemblies, mainly because dipcall’s default settings exclude variant-dense regions. Relative to PAV, longcallD exhibited higher FNRs but comparable FDRs on HiFi data and overall comparable accuracy on ONT data. Increasing sequencing depth improved recall but not precision for all callers, with longcallD’s F1 score plateauing beyond ∼30× coverage (**Supplementary Fig. 3**).

### Accurate haplotype reconstruction in tandem repeat regions

For more precise SV evaluation at the haplotype level, we applied aardvark^29^ to compute sequence-based genotype- and basepair-level F1 scores. Aardvark jointly considers small and structural variants. However, except for longcallD, dipcall, and PAV, other SV callers do not output small variants, which would lead to inherent bias in aardvark-based evaluation. For these SV-only callers, SNPs and small indels (<30 bp) from DeepVariant were merged with SV calls for fair comparison. Although this solution is not ideal due to different variant representations, it is the best we could do with these long-read SV callers.

Across both metrics, longcallD consistently achieved higher F1 scores than other long-read SV callers, particularly in tandem repeats (**Fig. 4a,b**). Compared with assembly-based callers, longcallD exhibited accuracy comparable to dipcall on HiFi data and higher accuracy on ONT data. PAV consistently achieved higher basepair F1 scores but lower genotype F1 scores than longcallD, reflecting differences in sequence accuracy versus allelic representation. Overall, longcallD, dipcall, and PAV achieved similar accuracy on both HiFi and ONT datasets and substantially outperformed other SV callers.

**Fig. 4.**
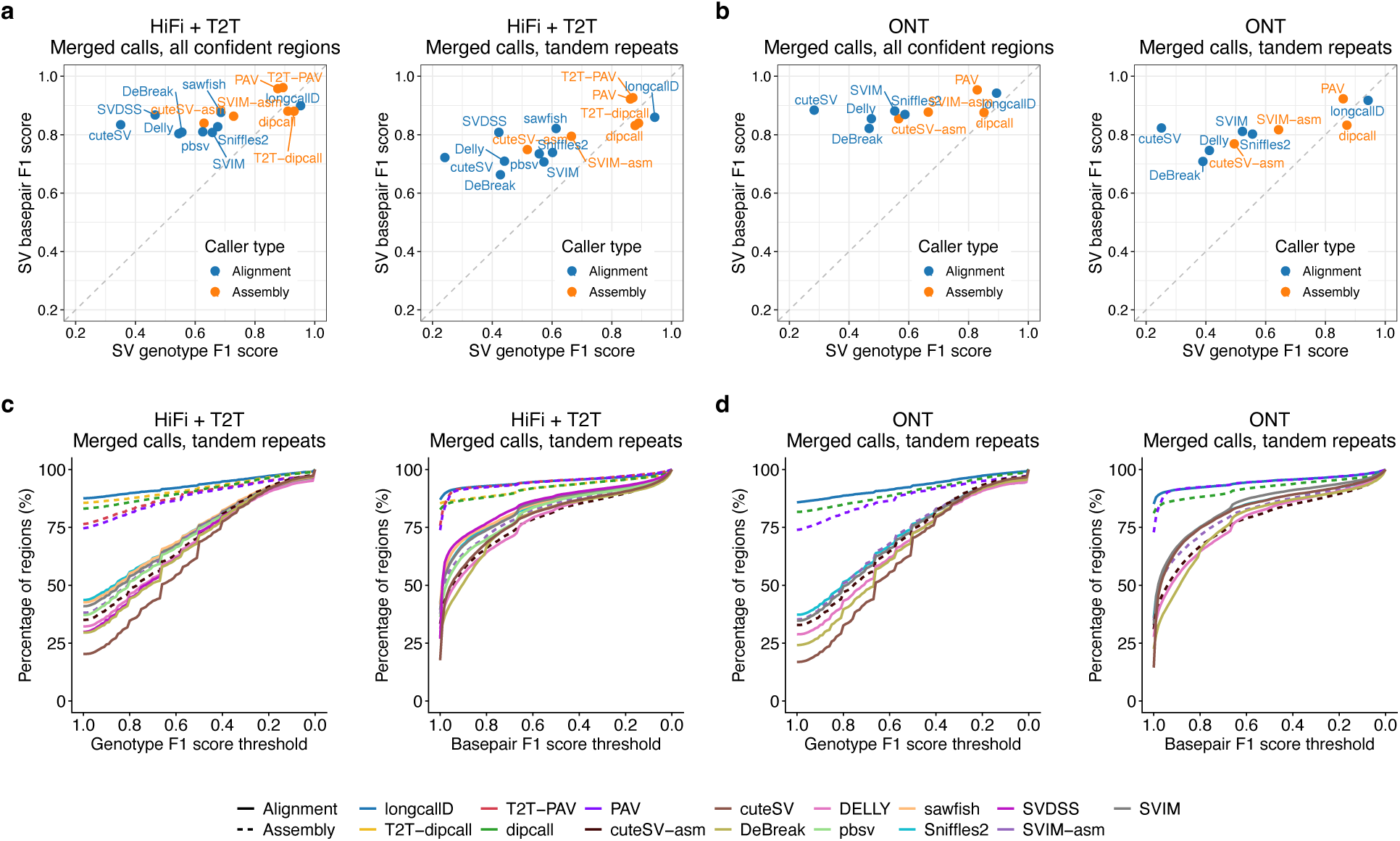
SV- and region-wise genotype and basepair F1 scores. **a**, SV-wise F1 scores in whole confident and tandem repeat regions on HiFi and T2T data, measured by aardvark. Dipcall, PAV, cuteSV-asm, and SVIM-asm were applied to long-read assembly alignments, T2T-dipcall and T2T-PAV were applied to T2T assembly alignments, all other callers were applied to long-read alignments. **b**, SV-wise F1 scores on ONT data. **c**, Region-wise F1 score distribution in tandem repeats on HiFi and T2T data, merged calls are shown for SV-only callers. Curves represent the fraction of tandem repeat regions (y-axis) achieving an F1 score above a given threshold (x-axis). **d**, Region-wise F1 score distribution in tandem repeats on ONT data.

To further characterize performance in tandem repeat regions, we examined region-wise F1 scores computed by aardvark for tandem repeat regions containing variant sequences of ≥ 50 bp. Region-wise F1 scores were calculated using all small and structural variants within each region, providing an aggregate measure of haplotype-level concordance between variant calls and the ground truth. For HiFi data, longcallD achieved perfect genotype and basepair F1 scores of 1.0 in 80.8% (13,294 of 16,452) and 81.9% (13,471 of 16,452) of regions, respectively. Similar fractions were observed for ONT data (81.2% for genotype and 81.9% for basepair). Across all genotype- and basepair-level F1 thresholds, longcallD consistently yielded a higher proportion of high-scoring regions than other long-read SV callers (Fig. 4c,d). Although PAV and dipcall exhibited slightly lower proportions of perfectly concordant regions, dipcall shows the highest precisions and PAV shows the highest recalls across most of the thresholds (**Supplementary Fig. 4**). Overall, the performance of longcallD, PAV, and dipcall substantially exceeded that of other long-read SV calling methods.

We also examined region-wise accuracy using the original variant calls from long-read SV callers (**Supplementary Fig. 5**). The basepair-level metrics of aardvark evaluate similarity at the base level, leading to high scores for highly similar sequences, whereas genotype-level metrics require exact matches for variants within the region. As expected, not jointly calling the small variants in the regions, all the other long-read SV callers have noticeably lower recalls, as they do not report nearby small variants. For precisions, they exhibited higher basepair-level precisions than genotype-level precisions, indicating the sequences of called variants are highly similar to the ground truth but not enough for haplotype reconstruction.

These results indicate that jointly calling small and large variants is critical for accurate haplotype reconstruction in repetitive genomic contexts. **Fig. 5** shows a representative tandem-repeat region on chromosome 5 in which longcallD correctly resolved three large insertions on both haplotypes, despite extensive clipping and fragmented, inconsistent alignments in the raw data. Through haplotype-aware alignment refinement, longcallD produced internally consistent haplotype-specific representations that exactly matched the ground truth. Across all evaluated datasets, longcallD, PAV, and dipcall achieved perfect genotype- and basepair-level F1 scores for this region, whereas SV-only callers generally yielded F1 scores below 0.80. An exception was SVDSS, which achieved a high basepair F1 score (0.99) but a substantially lower genotype F1 score (0.50) on the HiFi dataset.

**Fig. 5.**
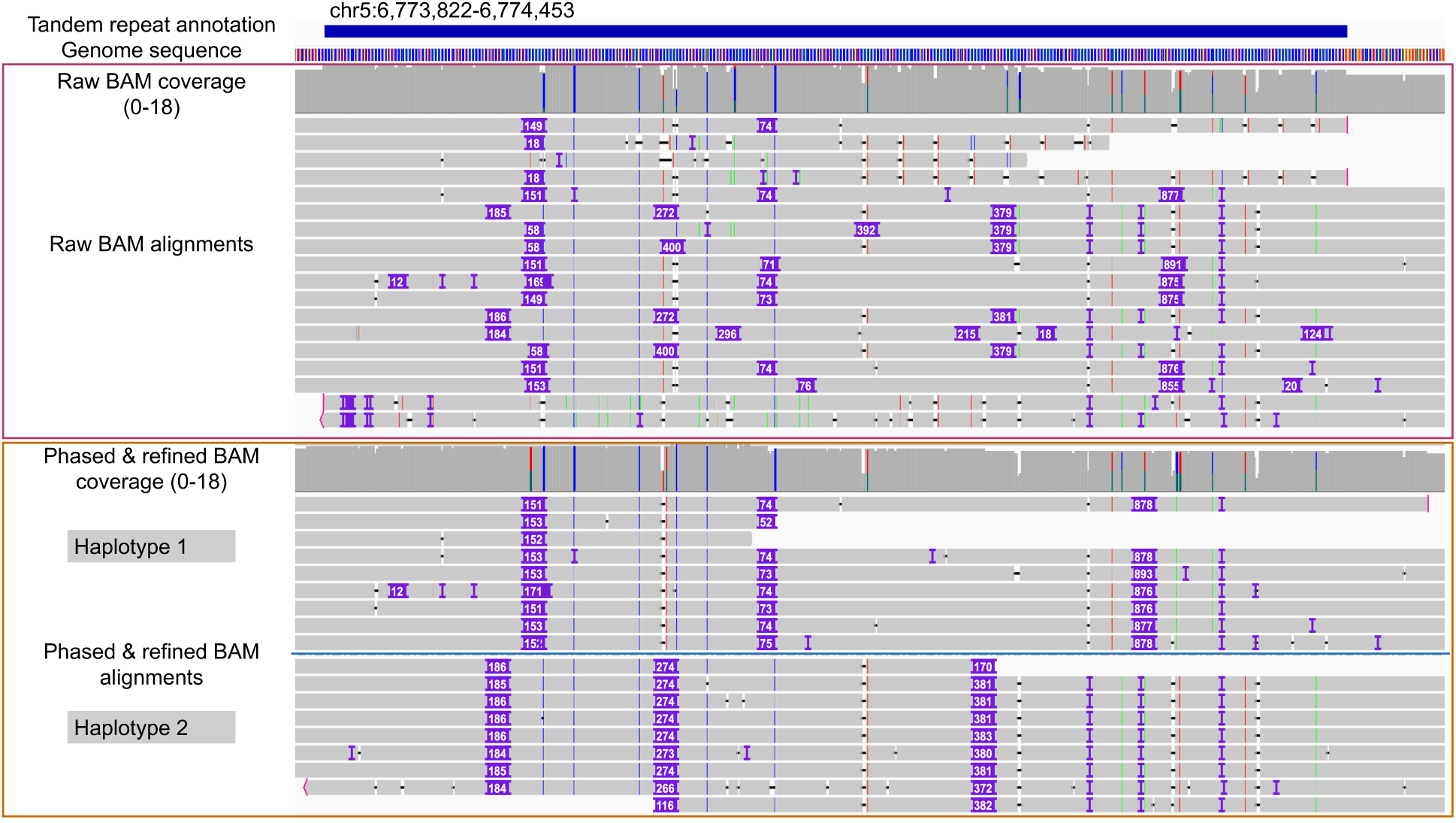
Haplotype-aware alignment refinement by longcallD. Top, tandem repeat annotation and reference genome sequence. Middle, HG002 raw HiFi long-read alignments displaying fragmented, inconsistent insertions and clippings. Bottom, haplotype-phased and refined alignments produced by longcallD. In this example, longcallD identified three large insertions on both haplotypes, together with other small variants.

### Mosaic SNV detection

We next evaluated longcallD’s performance in detecting mosaic SNVs of low VAF. DeepSomatic^31^ (tumor-only mode) and Clair-Mosaic^32^ were included for comparison. We first analyzed a COLO829 (melanoma) and COLO829BL (matched normal) mixture generated by the SMaHT network^33^ at a low tumor:normal ratio of 1:49. This design preserves the original germline haplotypes while introducing COLO829-specific somatic mutations at expected VAFs of ∼1% (heterozygous) or ∼2% (homozygous). We focused on a high-confidence reference set of 28,603 mosaic SNVs located in “easy” regions defined by the 1000 Genomes strict mask^5^. These truth SNVs were produced by SMaHT from high-coverage tumor-normal pairs.

On a 110× HiFi dataset, longcallD achieved the highest precision (52.0%) and recall (33.1%), outperforming DeepSomatic (47.6% precision, 20.7% recall) and Clair-Mosaic (17.2% precision, 3.4% recall) (**Fig. 6a**). LongcallD uniquely identified 4,343 true positive SNVs not detected by the other callers, the majority of which were supported by only two alternative reads (**Fig. 6b**). The alternative allele depth distribution of longcallD-specific calls was significantly lower than that of calls reported by DeepSomatic or Clair-Mosaic (P < 2.2 × 10^-16^), indicating superior sensitivity for variants with minimal read support. The relatively low recall observed for all methods is expected, as many mosaic SNVs with VAFs of 1–2% are not sampled even at 110× coverage under binomial sampling constraints. On an 87-fold ONT dataset of the same COLO829 mixture sample, while longcallD and DeepSomatic show a similar recall (15.3% and 15.8%) which are both lower than that on HiFi dataset, longcallD shows a superior precision of 39.0% compared to 5.6% for DeepSomatic (**Supplementary Fig. 6**).

**Fig. 6.**
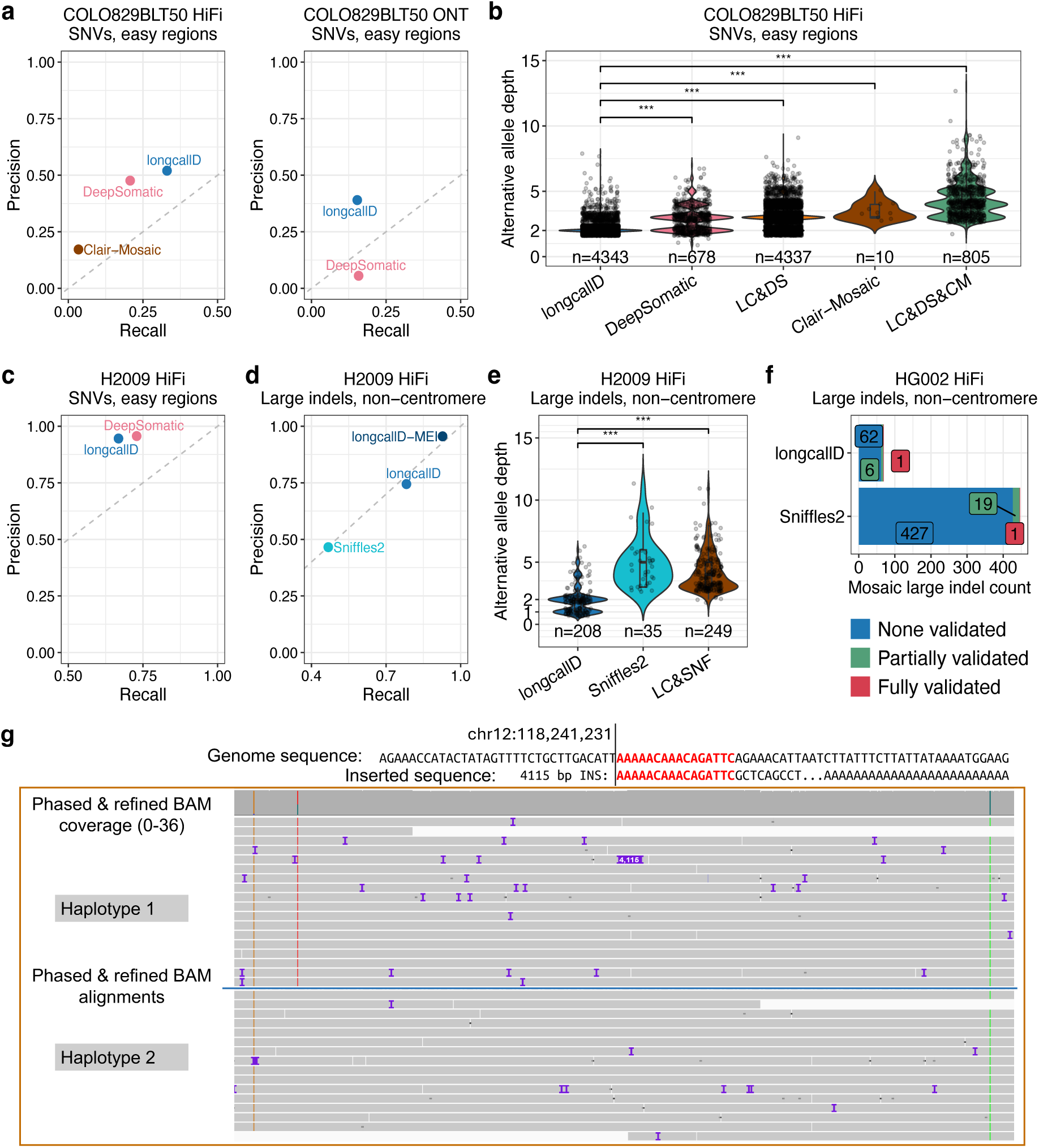
Low-VAF mosaic SNV and MEI calling. **a**, Precision and recall of mosaic SNV calls on COLO829BLT50 in easy regions using HiFi and ONT data. Clair-Mosaic did not report any mosaic SNVs on ONT dataset. **b**, Alternative allele depth of TP mosaic SNV calls on COLO829BLT50 using HiFi data. LongcallD, detected exclusively by longcallD; DeepSomatic, detected exclusively by DeepSomatic; LC&DS, detected by both longcallD and DeepSomatic but not by Clair-Mosaic; Clair-Mosaic, detected exclusively by Clair-Mosaic; LC&DS&CM, detected by all three callers. Wilcoxon rank-sum tests were used to compare the alternative allele depth distributions of longcallD-exclusive TP calls with those of other call sets (***P < 2.2 × 10⁻¹⁶). **c**, Precision and recall of mosaic SNV calls on H2009 in easy regions using synthetic HiFi data with a 1:10 tumor-normal mix ratio. Clair-Mosaic was excluded as it did not report any mosaic SNVs on this dataset. **d**, Precision and recall of mosaic large indel calls on H2009 using synthetic HiFi data in non−centromeric regions. longcallD-MEI denotes mosaic insertions retaining canonical mobile element features, including TSD and poly(A) tail sequences. **e**, Alternative allele depth of TP mosaic large indel calls on H2009. LongcallD, detect exclusively by longcallD; Sniffles2, detected exclusively by Sniffles2; LC&SNF: detected by both callers. **f**, Number of detected mosaic large indels that have no, some (≥1 but not all), and all supporting reads with indel in HG002 T2T alignments. **g**, Representative mosaic SVA insertion identified from NA12890 HiFi data. Top, 16-bp TSD, the corresponding reference sequence around the insertion site, and 73-bp poly(A) tail at the end of the insertion sequence. Bottom, long-read alignment phased by longcallD.

To assess performance at moderate VAFs, we generated a synthetic 80× HiFi dataset by mixing H2009 (lung cancer) and H2009BL (matched normal) cell lines at a tumor:normal ratio of 1:10. A published high-confidence somatic SNV call set comprising 140,043 variants supported by multiple sequencing technologies^28^ served as the ground truth. Clair-Mosaic was excluded from this analysis because it failed to report any mosaic SNVs on this dataset. Both longcallD and DeepSomatic achieved substantially higher precision (94.6% and 95.7%, respectively; **Fig. 6c**) than in the 1:49 mixture. However, longcallD showed slightly lower recall (66.8%) than DeepSomatic (72.9%), indicating that DeepSomatic is more sensitive for mosaic SNVs with moderate VAFs and stronger read support.

### Mosaic large indel detection

We next evaluated mosaic large indel detection using the synthetic H2009 dataset, with Sniffles2 (mosaic mode) included for comparison. As no curated ground-truth large-indel set is available for this dataset, we constructed an integrated reference set from four somatic SV callers including Severus^18^, SAVANA^34^, nanomonsv^35^, and minisv^36^, and applied to both tumor and matched normal samples (Methods). After excluding indels <100 bp, >50 kb or located in centromeric regions, 661 large indels detected by at least one caller were used to estimate precision, and 509 indels detected by all four callers were used to estimate recall.

Although longcallD (614) and Sniffles2 (610) reported similar numbers of mosaic large indels (≥100 bp), longcallD achieved substantially higher recall (78.2% versus 46.8%) (**Fig. 6d**). Whereas Sniffles2 requires at least three supporting reads for mosaic SV detection, longcallD accurately identified true mosaic indels supported by only one or two reads (**Fig. 6e**), while maintaining higher overall precision (74.4% versus 46.6%). These results indicate that haplotype-aware filtering enables sensitive detection of low-VAF mosaic indels without a corresponding increase in false positives.

We further constructed a mosaic MEI reference set using minisv on the tumor and normal H2009 samples. Insertions exhibiting canonical MEI features, i.e., TSD ≥2 bp and poly(A) tail ≥10 bp, and supported by at least one read in the synthetic HiFi dataset were retained, yielding 467 mosaic MEIs as ground truth. Applying the same TSD and poly(A) criteria to longcallD calls, we observed a precision of 95.5% and recall of 92.7%, demonstrating that explicit modeling of MEI sequence features markedly improves detection accuracy for low-VAF insertions (Fig. 6d).

As a negative control, we called mosaic SVs from HG002 HiFi reads and remapped all supporting reads to the HG002 T2T Q100 v1.1 assemblies. Supporting reads were aligned separately to the two T2T haplotypes and considered validated only if both haplotype alignments contained the corresponding indel. Mosaic SVs were classified as none validated (no supporting reads validated), partially validated (≥1 but not all supporting reads validated), or fully validated (all supporting reads validated). In total, 69 and 447 mosaic large indels (≥100 bp, <50 kb, non-centromeric) were detected by longcallD and Sniffles2, respectively. Among these, only a single mosaic deletion of 1474 bp (**Supplementary Fig. 7**) with five supporting reads was fully validated, and it was detected by both callers (**Fig. 6f**). Beyond this, most events were not validated in the T2T assemblies, considered as false positives. LongcallD reported substantially fewer such false positive calls, supporting its lower false-positive rate in mosaic SV detection. We suspect the longcallD false positives are caused by false mappings against the reference genome due to copy-number variants longer than reads.

**Fig. 6g** shows a representative mosaic SVA insertion on chromosome 12 detected in NA12890, supported by a single HiFi read. The insertion contains a 16-bp TSD and a 73-bp poly(A) tail, and its internal sequence matches known SVA elements. Because longcallD jointly calls germline and mosaic variants, it leverages haplotype phasing to confirm that the supporting read is consistent with other reads assigned to the same haplotype, thereby reducing the likelihood that the event reflects a sequencing artifact.

## Discussion

We present longcallD, a unified framework for jointly calling and phasing small variants, structural variants and low-VAF mosaic mutations from long-read sequencing data. Across HiFi and ONT datasets benchmarked against the HG002 T2T Q100 v1.1 truth set, longcallD achieves competitive small-variant accuracy and the strongest performance among alignment-based SV callers in tandem repeats, where variant representation and phasing are most error-prone.

A central design principle of longcallD is the explicit separation of the genome into clean and noisy regions. Rather than applying a uniform probabilistic model genome-wide, longcallD performs direct read-count–based calling in well-behaved regions and haplotype-aware MSA–based consensus reconstruction in low-complexity and breakpoint-associated regions. This targeted strategy addresses the non-uniform error landscape of long-read data and substantially improves SV detection in tandem repeats. Haplotype-level benchmarking with aardvark further demonstrates that accurate reconstruction of repeat loci requires joint modeling of small and large variants. SV-only approaches frequently produce internally inconsistent haplotypes, even when supplemented with small variants from external callers. In contrast, longcallD, PAV and dipcall, which natively call both variant classes, achieve markedly higher region-wise genotype concordance.

Compared to assembly-based approaches, longcallD attains broadly comparable haplotype-level accuracy while operating directly on read alignments. Using 16 threads on a standard computational server, longcallD completed analysis of a 40x HiFi alignment file in 19 minutes with less than 20GB peak memory, compared with 218 minutes and 40GB peak memory for a pipeline comprising DeepVariant, HiPhase and Sniffles2 in phase mode for calling phased small variants and SVs. De novo assembly-based frameworks typically demand several-fold greater runtime and memory. The design of longcallD eliminates the computational overhead and assembly-specific biases associated with de novo assembly, particularly in homopolymer-rich contexts on ONT data. Assembly-based methods retain advantages in more complex repeat structures, such as segmental duplications, but longcallD provides a practical alternative for routine analysis at standard whole-genome coverage.

LongcallD further extends haplotype-aware modeling to detect mosaic variants of low VAF, which are obscured or absent in assembled contigs. By requiring haplotype consistency among supporting reads, longcallD improves discrimination between true low-VAF mutations and sequencing artifacts. In tumor–normal mixture experiments at ∼1–2% VAF, longcallD achieved the highest combined precision and recall for mosaic SNVs and substantially outperformed existing tools for mosaic large indels, detecting events supported by as few as one or two reads without a commensurate increase in false positives. These results underscore the value of integrating germline phasing information into somatic variant inference.

Several limitations of longcallD remain. Homopolymer indel calling, especially on ONT data, is not as accurate as deep-learning-based methods. Complex SV events such as inversions and translocations which are more often to occur in tumor samples are not yet supported. Mosaic small indels are not currently reported due to their overlap with long-read error modes. Future work will extend variant classes, incorporate learned error models to improve homopolymer indel calling, and assess performance across diverse human genomes and tumor contexts.

As long-read sequencing becomes increasingly adopted in research and clinical genomics, unified frameworks that jointly resolve the full spectrum of variation with haplotype context will be essential. LongcallD provides an integrated solution for germline and mosaic variant analysis from long-read data, particularly in repetitive regions where accurate haplotype reconstruction is critical.

## Methods

### Identification of variant-calling noisy regions

#### Collecting candidate noisy regions

LongcallD processes all primary alignments with mapping quality ≥30. All bases with quality scores ≥10 that differ from the reference, including substitutions, insertions, and deletions extracted from CIGAR strings, are treated as candidate variants. LongcallD identifies noisy regions and records them as reference-coordinate intervals, each associated with a putative variant size *V*, which is later used for region merging. Four types of noisy regions are defined:

1. High-density difference windows. For each primary alignment, longcallD applies an extensible sliding window to count the total number of differing bases between the read and the reference. A window with length ≥*w* bp (100 for HiFi reads and 25 for ONT reads) and containing more than five differing bases is classified as a noisy region. The putative variant size *V* is defined as the total number of differing bases within the window.
2. Variants in low-complexity regions (LCRs). Any candidate variant overlapping an LCR is classified as noisy. LCRs are identified by applying the sdust algorithm^37^ with parameters “*W*=20, *T*=5” to the reference sequence. A noisy region is defined as the variant interval itself, and the putative variant size *V* is set to the number of differing bases in the variant.
3. Overlapping candidate variants. If two candidate variants overlap on the reference genome, their union interval is designated as a noisy region. The putative variant size *V* is defined as the larger number of differing bases between the overlapping variants. For clusters of more than two overlapping variants, this procedure is applied iteratively until no further overlaps remain.
4. Long-clipping-associated regions. For any alignment containing a clipped segment ≥30 bp, the 100 bp window flanking the clipped end is designated as a noisy region. For these regions, the putative variant size is fixed at *V* = 1.

#### Filtering of noisy regions

Identified noisy regions are filtered based on read-level support. Regions are retained only if (i) the total read coverage is ≥5; (ii) the number of alternate reads, i.e., supporting a variant or containing clippings, is ≥2; (iii) the ratio of alternate to total reads is ≥0.20.

#### Merging of noisy regions

After filtering, each remaining noisy region is first extended to fully cover any overlapping low-complexity regions. LongcallD then merges two adjacent noisy regions *i* and *j* if the distance between their intervals is ≤*V_min_*, where *V_min_* = min(*V_i_*, *V_j_*). The merged region is assigned a putative variant size *V_new_* = max(*V_i_*, *V_j_*). This merging procedure is applied recursively until no additional regions satisfy the merging criterion. Finally, each remaining noisy region is extended by 10 bp on both flanks.

### Clean-region germline variant calling and read haplotagging

Candidate variants located outside noisy regions are designated as clean-region variants. Variants with total coverage <5, alternate read coverage <2 or alternate allele ratio <0.20 are filtered out. Variants with alternate allele ratios between 0.20 and 0.80 are classified as heterozygous, whereas those with ratios >0.80 are classified as homozygous.

LongcallD phases variants and long reads in clean regions using an iterative haplotype extension strategy (**Fig. 1d**). A binary read–variant compatibility matrix is constructed, where entries indicate whether a read matches the reference allele (0), variant allele (1), or does not cover the site (-). Phasing is initialized from the heterozygous variant with the highest coverage, assigning the alternate and reference alleles to haplotypes 1 and 2, respectively. Reads covering this variant are assigned haplotypes accordingly. Phasing is then extended outward from the initial variant, first toward the left and subsequently toward the right along the genome.

Phasing proceeds iteratively by alternating between variant haplotype assignment and read haplotype assignment. Variants inherit haplotypes from overlapping reads by majority voting and will remain unassigned in the event of a tie. Conversely, reads inherit haplotypes from covered variants by majority voting and likewise remain unassigned if no majority exists. Iteration continues until haplotype assignments converge or until all reads and variants have been traversed ten times. Any two heterozygous variants covered by at least two shared reads are assigned to the same phase set. Each phase set is labeled by the smallest genomic coordinate among the variants it contains. Each read is assigned with the smallest phase set among all the variants it covers.

### Germline variant calling in noisy regions

#### Haplotype-aware consensus calling

Reads overlapping a noisy region are classified as fully spanning if they cover both region boundaries, or partially spanning if they cover only one boundary. For each noisy region, longcallD searches for a phase set satisfying: (i) at least one fully spanning read per haplotype; and (ii) at least two total reads per haplotype. If multiple phase sets meet these criteria, the set with the largest number of reads is selected; ties are resolved by choosing the left-most phase set.

Sequences corresponding to the noisy region are extracted from both fully and partially spanning reads and aligned separately for each haplotype using abPOA^38^ to generate haplotype-specific consensus sequences. Sequences are incorporated into the partial order graph in descending order of length, with fully spanning sequences added prior to partially spanning sequences. In practice, partially spanning sequences are first aligned to the longest fully spanning sequence using WFA^39^; the resulting alignment determines their start and end nodes in the graph, enabling end-to-end sequence-to-graph alignment.

#### Haplotype-agnostic consensus calling

For noisy regions lacking a valid phase set, regions with <5 fully spanning reads are skipped. Otherwise, haplotype-agnostic consensus calling is performed using fully spanning reads only. Sequences are sorted by length and aligned using abPOA, and the resulting partial order graph is converted into a row–column multiple sequence alignment.

Alignment columns are designated as candidate heterozygous columns if two bases each have coverage ≥2 and ≤0.8×*Cov_total_*, where *Cov_total_* denotes the total column coverage. Columns are classified as substitutions when both bases are non-gaps and as indels when one base is a gap. Based on these columns, abPOA computes a pairwise distance matrix among sequences, weighting substitutions as 2 and indels as 1. K-medoids clustering is applied to partition sequences into two clusters. Clustering is accepted if each cluster contains 20–80% of sequences. If accepted, two haplotype-specific consensus sequences are generated; otherwise, a single consensus sequence is produced, indicating a homozygous noisy region.

#### Variant calling from haplotype consensus sequences

Consensus sequences are aligned to the reference sequence of the noisy region using WFA. For homozygous noisy regions, variants are called directly from the alignment. For heterozygous regions, variants are called independently from each haplotype consensus sequence. Variants present in both haplotypes are reported as homozygous, whereas haplotype-specific variants are reported as heterozygous.

#### Haplotype phasing in noisy regions

After variant calling across all noisy regions, longcallD performs a second round of haplotype phasing jointly across clean-region and noisy-region variants. The procedure mirrors clean-region phasing, except that reads inherit haplotypes from covered variants using weighted majority voting: clean-region variants are assigned weight 2 and noisy-region variants weight 1. For noisy-region variants, the read–variant compatibility matrix is constructed by aligning haplotype consensus sequences to individual reads to determine allele support.

### Low-VAF mosaic variant calling

LongcallD performs additional low-VAF mosaic variant calling after germline variant calling. Currently, mosaic SNVs and large indels are supported, whereas mosaic small indels are not called due to the difficulty of distinguishing them from long-read sequencing errors.

#### Collecting candidate mosaic variants

During germline variant calling, filtered SNPs and large indels (length ≥30 bp) with alternate allele ratio <0.20 and alternate read coverage ≥2 are recorded as candidate mosaic variants. Candidate mosaic MEIs are identified by requiring: (i) the presence of TSD ≥2 bp and poly(A) tail sequences ≥10 bp within large insertions; and (ii) the inserted sequence contains ≥3 non-overlapping 15-bp subsequences of any known mobile elements (e.g., Alu, L1, and SVA). For mosaic MEIs, the alternate read coverage threshold is reduced to one.

#### Selection of phase set

For each candidate mosaic variant, longcallD collects all reference and alternate reads covering the variant and identifies a phase set satisfying the following criteria: (i) it contains the largest number of alternate reads among all phase sets; (ii) all alternate reads originate from the same haplotype; (iii) the number of alternate reads does not exceed the number of reference reads from the same haplotype and phase set; and (iv) the total number of reference and alternate reads is ≥5. Once a phase set was selected, only alternate reads from this phase set will be used in the following filtering step.

#### Filtering candidate mosaic variants

Candidate mosaic SNVs are filtered out if any of the following conditions are met: (i) the distance to any germline variant is <5 bp; (ii) the median base quality of alternate bases is lower than the median base quality of all bases; (iii) the median of the minimum base quality scores within 7-bp windows (variant base ± 3 bp) around alternate bases is lower than the first quartile base quality of all bases; (iv) the median distance between alternate bases and the nearest indel sequencing error is <5 bp; (v) ≥50% of alternate bases lie within 500 bp of a dense-difference window (50 bp window containing >5 differing bases); (vi) ≥ 50% of alternate bases lie within 100 bp of a clipping event with length ≥100 bp; or (vii) ≥ 50% of alternate reads contain sequencing errors within LCRs covering the variant.

Candidate mosaic large indels are filtered out if any of the following conditions are met: (i) the variant lies within 500 bp of a germline indel and either ≥80% of the differing sequence is low-complexity or the sequence similarity between the two variants is ≥80%; (ii) the alternate read count equals 1 and ≥80% of the variant sequence is low-complexity; (iii) ≥50% of alternate bases lie within 500 bp of a dense-difference window (50 bp window containing >5 differing bases); or (iv) ≥50% of alternate bases lie within 100 bp of a clipping event with length ≥100 bp.

#### Post-filtering of mosaic variants

Candidate mosaic variants located within 1 kb windows containing more than five mosaic variants are removed. supporting filtered mosaic variants are marked as invalid for mosaic calling. Remaining mosaic variants supported by any invalid reads are subsequently filtered out.

### ONT-specific processing

Due to the higher error rate and increased prevalence of artifacts in ONT reads compared with PacBio HiFi reads, longcallD applies additional ONT-specific filters.

#### Palindromic clippings

If a supplementary alignment indicates that a clipped sequence maps to the same locus as the primary alignment and the overlapping region covers ≥ 90% of the primary alignment, the clipping is classified as palindromic and is not treated as a noisy region.

#### Stranded bias

For each candidate germline variant, longcallD performs a Fisher’s exact test comparing the observed numbers of forward- and reverse-strand alternate reads with the expected counts (assumed equal). Variants with P < 0.01 are classified as strand-biased and filtered out. For mosaic variant calling, in addition to the criteria above, candidate mosaic SNVs are filtered out if all alternate reads originate exclusively from either the forward or the reverse strand.

### Harmonizing somatic SV call set as ground-truth mosaic variants

A de novo assembly of HiFi reads from the H2009 normal cell line was generated using hifiasm (v0.19.6). HiFi long reads from the H2009 tumor sample were aligned to four references: GRCh38, CHM13v2 linear genome^40^, and the normal-sample de novo assembly using minimap2 (v2.28)^41^, and to CHM13 graph genome^42^ using minigraph (v0.21)^43^. Small variants in the normal cell line were called from long-read data using Clair3^7^ with phasing enabled. The resulting variant calls and read alignments were provided to WhatsHap^44^ to generate the phased alignment files.

We applied minisv (v0.1.2)^36^ in tumor–normal paired mode, integrating the tumor alignments described above together with GRCh38-based alignments of the normal sample to identify somatic SVs, with coordinates reported on GRCh38. An initial GRCh38-based SV call set was generated using parameters “-c 1 -s 0” to retain a comprehensive set of candidates where all calls have at least one supporting read, which was used for mosaic MEI calling evaluation. For downstream consensus harmonization, a more stringent cutoff, i.e., “-c 2 -s 0” was applied to only retain calls with at least two supporting reads.

In parallel, somatic SVs were identified on GRCh38 using tumor–normal paired mode in Severus (v1.0)^18^ and SAVANA (v1.2)^34^ with the phased alignment file, enabling VNTR-aware mode under default settings. Nanomonsv (v0.7)^35^ was applied to the original alignment file to generate GRCh38-based somatic SV calls. For all methods, read ID output was enabled. Each SV call set, together with its corresponding read ID file, was processed using minisv filterasm with breakpoint signals derived from de novo assembly–based read alignments. This step restricted calls to high-confidence regions (https://github.com/lh3/minisv/blob/master/data/hs38.reg.bed), excluding pseudo-autosomal regions on chromosomes X and Y, centromeric regions, and acrocentric short arms. Finally, all filtered SV call sets were harmonized using the minisv union with parameters “-p -c 5”, which identifies consensus SVs across methods and retains calls supported by at least five reads. The resulting consensus set was used as the reference mosaic SV call set for evaluation.

## Data availability

Human reference genome: GRCh38: ftp://ftp.ncbi.nlm.nih.gov/genomes/all/GCA/000/001/405/GCA_000001405.15_GRCh38/seqs_for_alignment_pipelines.ucsc_ids/GCA_000001405.15_GRCh38_no_alt_analysis_set.fna.gz, CHM13v2: https://s3-us-west-2.amazonaws.com/human-pangenomics/T2T/CHM13/assemblies/analysis_set/chm13v2.0.fa.gz, HG002 Q100: https://s3-us-west-2.amazonaws.com/human-pangenomics/T2T/HG002/assemblies/hg002v1.1.fasta.gz, and CHM13 graph genome: https://zenodo.org/records/16728828/files/CHM13-464.gfa.gz?download=1; Alignments of HG002 PacBio HiFi reads: s3://human-pangenomics/submissions/40399FDD-59DE-43D1-B3A3-DFF0C6E64FAC--YALE_VARIANT_CALLS_R2/samples/HG002/alignments/GRCh38_no_alt/ hifi_reads/HG002.GRCh38_no_alt.sorted.bam; Alignments of HG002 ONT R10.4.1 super-accuracy (SUP) reads: s3://ont-open-data/giab_2025.01/basecalling/sup/HG002/PAW70337/calls.sorted.bam; Alignments of COLO829BLT50 PacBio HiFi reads: https://data.smaht.org/output-files/ef3ab86a-a00d-4cb8-bf3e-10447d41df07/; Alignments of COLO829BLT50 ONT R10.4.1 SUP reads: https://data.smaht.org/output-files/86b8f2a9-75f9-4e70-a1cc-6c8eb62ed09b/; Sequence Read Archive (SRA) accession id of H2009 adenocarcinoma cell line PacBio HiFi reads: SRR28305172; SRA accession id of H2009 B lymphoblast cell line PacBio HiFi reads: SRR28305169; Alignments of NA12890 PacBio HiFi reads: s3://platinum-pedigree-data/data/hifi/mapped/GRCh38/NA12890.GRCh38.haplotagged.bam.

## Code availability

LongcallD is released under the MIT license and is freely available at https://github.com/yangao07/longcallD.

## Notes

### Competing Interest Statement

The authors have declared no competing interest.

