## Supplementary Materials for "LongcallD: joint calling and phasing of small, structural and mosaic variants from long reads"

### Supplementary Information

### 1 Software versions and commands

#### 1.1 Germline variant calling

1.1.1 longcallD v0.0.10

HiFi reads:

```
longcallD call <ref.fa> <hifi.bam> -t <nThreads> --min-cov 6 --alt-cov 3 \
     --min-mapq 5 --min-sv-len 30 -o <output.vcf>
```

ONT reads:

```
longcallD call --ont <ref.fa> <ont.bam> -t <nThreads> --min-cov 6 \
     --alt-cov 3 --min-mapq 5 --min-sv-len 30 -o <output.vcf>
```

1.1.2 DeepVariant v1.9.0

HiFi reads:

```
singularity run -B <inputDir>,<outputDir> deepvariant.sif \
     /opt/deepvariant/bin/run_deepvariant --model_type PACBIO --ref <ref.fa> \
     --reads <hifi.bam> --num_shards <nThreads> --output_vcf <output.vcf>
```

ONT reads:

```
singularity run -B <inputDir>,<outputDir> deepvariant.sif \
     /opt/deepvariant/bin/run_deepvariant --model_type ONT_R104 --ref <ref.fa> \
     --reads <ont.bam> --num_shards <nThreads> --output_vcf <output.vcf>
```

1.1.3 Clair3 v1.2.0

HiFi reads:

```
singularity run -B <inputDir>,<outputDir> clair3.sif \
     /opt/bin/run_clair3.sh -f <ref.fa> -b <hifi.bam> -p hifi \
     --model_path=/opt/models/hifi_revio --min_coverage=6 -min_mq=5 \
     --use_longphase_for_intermediate_phasing \
     --use_longphase_for_final_output_phasing \
```

```
      --output=<outputDir>
```

ONT reads:

```
singularity run -B <inputDir>,<outputDir> clair3.sif \
      /opt/bin/run_clair3.sh -f <ref.fa> -b <ont.bam> -p ont \
      --model_path=/opt/models/r1041_e82_400bps_sup_v500 --min_coverage=6 \
      -min_mq=5 --use_longphase_for_intermediate_phasing \
      --use_longphase_for_final_output_phasing --output=<outputDir>
```

1.1.5 cuteSV v2.1.1

HiFi reads:

```
cuteSV --threads <nThreads> --min_mapq 5 --min_support 3 --min_size 30 \
      --genotype --max_cluster_bias_INS 1000 --diff_ratio_merging_INS 0.9 \
      --max_cluster_bias_DEL 1000 --diff_ratio_merging_DEL 0.5 \
      <hifi.bam> <ref.fa> <output.vcf> <outputDir>
```

ONT reads:

```
cuteSV --threads <nThreads> --min_mapq 5 --min_support 3 --min_size 30 \
      --genotype --max_cluster_bias_INS 100 --diff_ratio_merging_INS 0.3 \
      --max_cluster_bias_DEL 100 --diff_ratio_merging_DEL 0.3 \
      <ont.bam> <ref.fa> <output.vcf> <outputDir>
```

1.1.6 DeBreak v1.3

HiFi/ONT reads:

```
debreak --thread <nThreads> --min_size 30 --min_support 3 --min_quality 5 \
      --rescue_dup --rescue_large_ins --poa --ref <ref.fa> --bam <hifi/ont.bam> \
      --outpath <outputDir>
```

1.1.7 DELLY v1.3.2

HiFi reads:

```
delly lr --technology pb --mapqual 5 --min-clique-size 3 --geno-qual 5 \
      --genome <ref.fa> <hifi.bam> --outfile <output.vcf>
```

ONT reads:

```
delly lr --technology ont --mapqual 5 --min-clique-size 3 --geno-qual 5 \
      --genome <ref.fa> <hifi.bam> --outfile <output.vcf>
```

1.1.8 pbsv v2.10.0

HiFi reads:

```
pbsv discover --hifi --sample <sample> --min-mapq 5 --tandem-repeats \
      <trf_bed> <hifi.bam> <svsig.gz>
pbsv call --hifi --preserve-non-acgt --min-sv-length 30 --num-threads \
      <nThreads> <ref.fa> <svsig.gz> <output.vcf>
```

1.1.9 sawfish v0.12.8

HiFi reads:

```
sawfish discover --threads <nThreads> --min-indel-size 30 --min-sv-mapq 5 \
      --disable-path-canonicalization --ref <ref.fa> --bam <hifi.bam> \
      --output-dir <discoverOutputDir>
sawfish joint-call --threads <nThreads> --min-sv-mapq 5 \
      --sample <discoverOutputDir> --output-dir <callOutputDir>
```

1.1.10 Sniffles2 v2.5.3

HiFi/ONT reads:

```
sniffles --minsupport 3 --minsvlen 30 --mapq 5 --threads <nThreads> \
      --reference <ref.fa> --tandem-repeats <trf.bed> --input <hifi/ont.bam> \
      --vcf <output.vcf>
```

1.1.11 SVDSS v2.0.0

HiFi reads

```
SVDSS index –t <nThreads> -d <ref.fa> -o <ref.fmd>
SVDSS smooth --threads <nThreads> --min-mapq 5 --reference <ref.fa> \
      --bam <hifi.bam> > <smoothed.bam>
samtools index -@ <nThreads> <smoothed.bam>
SVDSS search --threads <nThreads> --index <ref.fmd> --bam <smoothed.bam> \
      > <specifics.txt>
SVDSS call --threads <nThreads> --min-cluster-weight 3 --min-sv-length 30 \
      --min-mapq 5 --reference <ref.fa> --bam <smoothed.bam> \
      --sfs <specifics.txt> > <output.vcf>
```

1.1.12 SVIM v2.0.0

HiFi/ONT reads:

```
svim alignment --min_mapq 5 --min_sv_size 30 --minimum_depth 3 \
      --interspersed_duplications_as_insertions <outputDir> \
      <hifi/ont.bam> <ref.fa>
```

1.1.13 dipcall v0.3

HiFi/ONT/T2T assemblies:

```
run-dipcall -t <nThreads> -x <ref.PAR.bed> -W <ref.k19.txt> \
      <outputPrefix> <ref.fa> <hap1.fa> <hap2.fa> > <outputPrefix>.mak
```

Here, “<ref.PAR.bed>” and “<ref.k19.txt>” are extracted from https://s3-us-west-2.amazonaws.com/human-pangenomics/submissions/40399FDD-59DE-43D1-B3A3-DFF0C6E64FAC--YALE_VARIANT_CALLS_R2/references/GRCh38_no_alt.tar.gz

1.1.14 PAV v2.4.6

HiFi/ONT/T2T assemblies:

```
cat << EOF > config.json
{
"reference": "<ref.fa>"
}
EOF
cat << EOF > assemblies.tsv
NAME\tHAP1\tHAP2
<sampleName>\t<hap1.fa>\t<hap2.fa>
EOF
/opt/pav/files/docker/run --notemp -c <nThreads>
```

1.1.15 cuteSV-asm v2.1.1

HiFi/ONT assemblies

```
cuteSV --threads <nThreads> --sample <sampleName> --min_mapq 5 \
      --min_support 1 --min_size 30 --max_size -1 --report_readid --genotype \
      --max_split_parts -1 --merge_del_threshold 500 --merge_ins_threshold 500 \
      --max_cluster_bias_INS 1000 --diff_ratio_merging_INS 0.9 \
      --max_cluster_bias_DEL 1000 --diff_ratio_merging_DEL 0.5 \
      <hap_merged.bam> <ref.fa> <asm.intermediate.vcf> <outputDir>
```

```
/opt/cutesv-asm/src/cuteSV/diploid_calling.py --hap1 '<sampleName>#1' \
     --hap2 '<sampleName>#2' <asm.intermediate.vcf> <output.vcf>
```

1.1.16 SVIM-asm v1.0.3

HiFi/ONT assemblies

```
svim-asm diploid --min_mapq 5 --min_sv_size 30 --query_names \
     --interspersed_duplications_as_insertions <outputDir> \
     <hap1.bam> <hap2.bam> <ref.fa>
```

#### 1.2 Low allele-fraction mosaic variant calling

1.2.1 longcallD v0.0.10

HiFi reads:

```
longcallD call -s <ref.fa> <hifi.bam> -t <nThreads> --min-sv-len 30 \t
     -T <AluY_L1_SVA_cons_noPA.fa> -o <output.vcf>
```

ONT reads:

```
longcallD call -s --ont <ref.fa> <hifi.bam> -t <nThreads> --min-sv-len 30 \
     -T <AluY_L1_SVA_cons_noPA.fa> -o <output.vcf>
```

“<AluY_L1_SVA_cons_noPA.fa>” can be downloaded from
https://github.com/yangao07/longcallD/blob/main/anno/AluY_L1_SVA_cons_noPA.fa

1.2.2 DeepSomatic v1.8.0

HiFi reads:

```
singularity run -B <inputDir>,<outputDir> deepsomatic.sif run_deepsomatic \
     -model_type=PACBIO_TUMOR_ONLY --ref=<ref.fa> --reads_tumor=<hifi.bam> \
     --output_vcf=<output.vcf> --num_shards=<nThreads> \
     --intermediate_results_dir=<intermediateDir> \
     --use_default_pon_filtering=true
```

ONT reads:

```
singularity run -B <inputDir>,<outputDir> deepsomatic.sif run_deepsomatic \
     -model_type=ONT_TUMOR_ONLY --ref=<ref.fa> --reads_tumor=<ont.bam> \
```

```
    --output_vcf=<output.vcf> --num_shards=<nThreads> \
    --intermediate_results_dir=<intermediateDir> \
    --use_default_pon_filtering=true
```

1.2.3 Clair-Mosaic v0.1.0

HiFi reads:

```
singularity run -B <inputDir>,<outputDir> <clair-mosaic.sif> \
    /opt/bin/run_clair_mosaic -bam_fn <hifi.bam> --ref_fn <ref.fa> \
    --threads <nThreads> --snv_min_af 0.01 --min_coverage 2 \
    --platform hifi_revio --conda_prefix /opt/micromamba/envs/clair-mosaic \
    --output_dir <outputDir>
```

ONT reads:

```
singularity run -B <inputDir>,<outputDir> <clair-mosaic.sif> \
    /opt/bin/run_clair_mosaic -bam_fn <ont.bam> --ref_fn <ref.fa> \
    --threads <nThreads> --snv_min_af 0.01 --min_coverage 2 \
    --platform ont_r10_dorado_sup_5khz \
    --conda_prefix /opt/micromamba/envs/clair-mosaic \
    --output_dir <outputDir>
```

1.2.4 Sniffles2 v2.5.3

HiFi/ONT reads:

```
sniffles --input <hifi/ont.bam> --mosaic --tandem-repeats <trf.bed> \
    --mosaic-af-min 0.01 --vcf <output.vcf> -t <nThreads>
```

"<trf.bed>" can bed downloaded from https://github.com/fritzsedlazeck/Sniffles/blob/master/annotations/human_GRCh38_no_alt_analysis_set.trf.bed

#### 1.3 Tumor-normal paired somatic variant calling

1.3.1 hifiasm v0.19.6

Normal sample:

```
hifiasm -t <nThreads> -o <normal_asm> <hifi.fq>
```

1.3.2 minimap2 v2.28

Normal sample:

```
minimap2 -x map-hifi -t <nThreads> <hg38.fa> <hifi.fq> > <normal_hg38.paf>
```

Tumor sample:

```
minimap2 -cx map-hifi -s50 -t <nThreads> <hg38.fa> <hifi.fq> > <tumor_hg38.paf>

minimap2 -cx map-hifi -s50 -t <nThreads> <chm13.fa> <hifi.fq> >
<tumor_chm13v2.paf>

minimap2 -cx map-hifi -s50 -I100g -t <nThreads> <normal_asm.fa> <hifi.fq> >
<tumor_asm.paf>
```

1.3.3 minigraph v0.21

Tumor sample:

```
minigraph -cxlr -t <nThreads> <chm13.gfa> <hifi.fq > > <tumor_graph.gaf>
```

1.3.4 minisv v0.1.2

```
minisv extract -n tumor <tumor_hg38.paf> > <tumor_hg38.rsv>

minisv extract -n tumor <tumor_chm13.paf> > <tumor_chm13.rsv>

minisv extract -n tumor -q0 -Q0 <tumor_asm.paf> > <tumor_asm.rsv>

minisv extract -x 5 -n tumor <tumor_graph.gaf> > <tumor_graph.rsv>

minisv extract -n normal <normal_hg38.paf> > <normal_hg38.rsv>

minisv isec <tumor_hg38.rsv> <tumor_chm13.rsv> | \

     minisv isec - < tumor_graph.rsv> | \

     minisv isec - <tumor_asm.rsv> > <tumor_isec.rsv>

cat <tumor_isec.rsv> <normal_hg38.rsv> | sort -k1,1 -k2,2 -S4g |\

     minisv merge -c1 -s0 - | grep tumor | grep -v normal > <output.msv>
```

For consensus harmonization:

```
cat <tumor_isec.rsv> <normal_hg38.rsv> | sort -k1,1 -k2,2 -S4g |\

     minisv merge -c2 -s0 - | grep tumor | grep -v normal > <output.msv>
```

1.3.5 Severus v1.0

```
singularity run -B <inputDir>,<outputDir> clair3.sif /opt/bin/run_clair3.sh \

     -f <ref.fa> -b <hifi.bam> -p hifi --model_path=/opt/models/hifi_revio \

     --output=<clair3_output> --enable_phasing --longphase_for_phasing
```

```
whatshap haplotag --reference <ref.fa> \
      <clair3_output>/phased_merge_output.vcf.gz <hifi.bam> -o <tag.bam> \
      --ignore-read-groups --tag-supplementary --skip-missing-contigs \
      --output-threads=<nThreads>
samtools index <tag.bam>
severus.py --target-bam <tumor_tag.bam> --control-bam <control_tag.bam> \
      --out-dir <output_folder> -t <nThreads> --vntr-bed <trf.bed> \
      --phasing-vcf <clair3_output>/phased_merge_output.vcf.gz --output-read-ids
```

“<trf.bed>” can bed downloaded from https://github.com/fritzsedlazeck/Sniffles/blob/master/annotations/human_GRCh38_no_alt_analysis_set.trf.bed

1.3.6 SAVANA v1.2

```
singularity run -B <inputDir>,<outputDir> clair3.sif /opt/bin/run_clair3.sh \
      -f <ref.fa> -b <tumor.bam> -p hifi --model_path=/opt/models/hifi_revio \
      --output=<tumor_output> --enable_phasing --longphase_for_phasing
singularity run -B <inputDir>,<outputDir> clair3.sif /opt/bin/run_clair3.sh \
      -f <ref.fa> -b <nomal.bam> -p hifi --model_path=/opt/models/hifi_revio \
      --output=<normal_output> --enable_phasing --longphase_for_phasing


whatshap haplotag --reference <ref.fa> \
      <clair3_output>/phased_merge_output.vcf.gz <tumor.bam> -o <tumor_tag.bam> \
      --ignore-read-groups --tag-supplementary --skip-missing-contigs \
      --output-threads=<nThreads>
whatshap haplotag --reference <ref.fa> \
      <clair3_output>/phased_merge_output.vcf.gz <normal.bam> \
      -o <normal_tag.bam> --ignore-read-groups --tag-supplementary \
      --skip-missing-contigs --output-threads=<nThreads>
samtools index <tumor_tag.bam>
samtools index <normal_tag.bam>
savana --threads 12 --tumour <tumor_tag.bam> --normal <control_tag.bam> \
      --outdir <output_folder> --ref <ref.fa> --pb --cn_binsize 10 --phased_vcf \
      <clair3_output>/phased_merge_output.vcf.gz --contigs <ref_contig.txt>
```

1.3.7 Nanomonsv v0.2

```
nanomonsv parse --reference_fasta <ref.fa> <normal.bam> <normal_prefix>

nanomonsv parse --reference_fasta <ref.fa> <tumor.bam> <tumor_prefix>

nanomonsv get <tumor_prefix> <tumor.bam> <ref.fa> \
     --control_prefix <normal_prefix> --control_bam <normal.bam>
```

#### 1.4 De novo genome assembly

1.4.1 HiFi reads: hifiasm v0.19.9

```
hifiasm -o <sampleASM> -t <nThreads> <hifi.fq>
```

1.4.2 ONT reads: hifiasm v0.25.0

```
hifiasm -ont -o <sampleASM> -t <nThreads> <ont.fq>
```

#### 1.5 Read and assemblies mapping

HG002 HiFi long reads were assembled using hifiasm-v0.19.9 with default settings. HG002 ONT long reads were assembled using the latest hifiasm-v0.25.0 with parameter "--ont". HiFi assembiles, HiFi raw reads, and ONT assemblies were aligned to GRCh38 human reference genome using Winnowmap (v2.03). ONT raw reads were aligned to GRCh38 human reference genome using minimap2 from Dorado-v0.8.2 package.

1.5.1 HiFi read mapping

```
meryl count k=15 <output> merylDB <ref.fa>

meryl print greater-than distinct=0.9998 merylDB > <repetitive_k15.txt>

winnowmap -ax map-pb -W <repetitive_k15.txt> <ref.fa> <hifi.fq> > <hifi.sam>
```

1.5.2 ONT read mapping

```
minimap2 -ax lr:hq <ref.fa> <ont.fq> > <ont.sam>
```

1.5.3 HiFi and ONT assemblies mapping:

```
meryl count k=19 output merylDB <asm.fa>

meryl print greater-than distinct=0.9998 merylDB > <repetitive_k19.txt>

winnowmap -W <repetitive_k19.txt> -ax asm20 <asm.fa> > <asm.sam>
```

#### 1.6 Variant calling benchmarking

1.6.1 hap.py v0.3.15

```
hap.py --pass-only --threads <nThreads> -r <ref.fa> \
     -f <small_benchmark.bed> -o <outputPrefix> <small_benchmark.vcf>
<caller.vcf>
```

1.6.2 Truvari v5.3.0

```
truvari bench -f <ref.fa> --includebed <sv_benchmark.bed> \
     -o <outputPrefix> -p 0.7 -P 0.7 -b <sv_benchmark.vcf> \
     -c <caller.vcf> --passonly --pick ac --dup-to-ins
truvari refine -m '--auto --thread <nThreads>' -f <ref.fa> \
     --regions <outputPrefix>/candidate.refine.bed \
     --use-original-vcfs <outputPrefix>
```

1.6.3 aardvark v0.9.0

```
aardvark compare -r <ref.fa> -t <benchmark.vcf> -q <caller.vcf> \
     -b <benchmark.bed> -o <outputPrefix> --min-variant-gap 1000 \
     --threads <nThreads>
```

#### 2 Supplementary figures

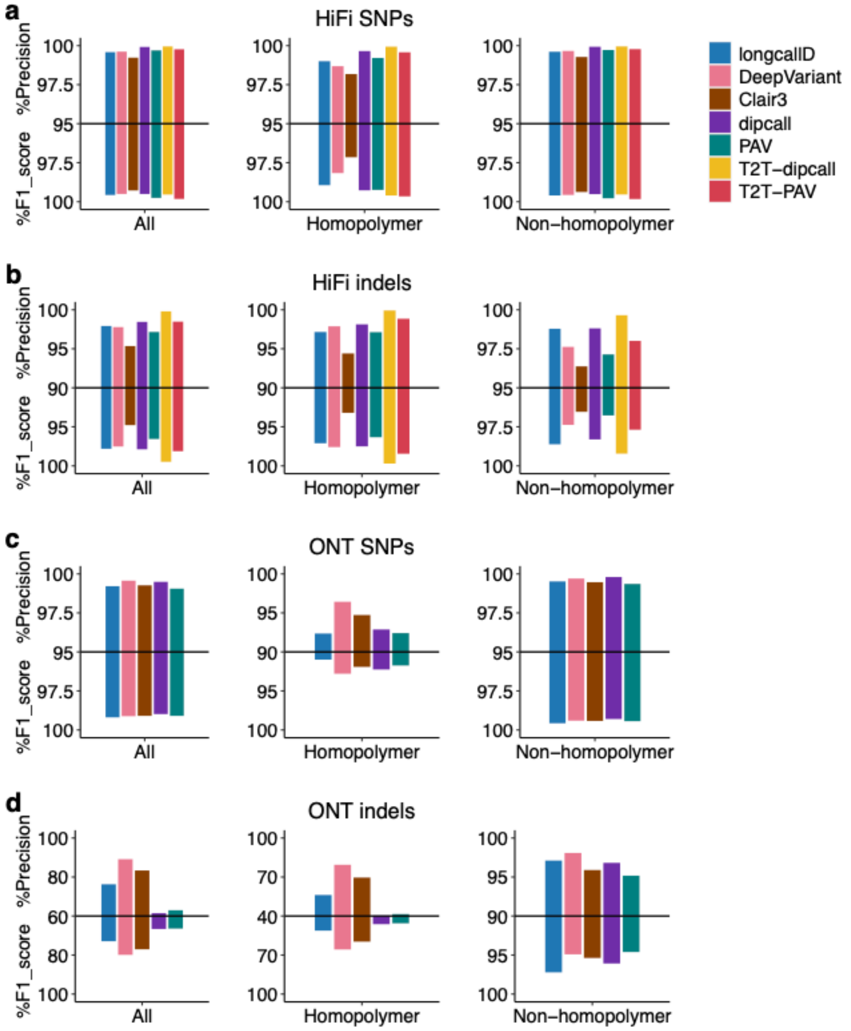


**Supplementary Fig. 1 | Small-variant calling precisions and F1 scores.** a, Precision and F1 scores for SNPs (HiFi and T2T). b, Precision and F1 scores for indels (HiFi and T2T). c, Precision and F1 scores for SNPs (ONT). d, Precision and F1 scores for indels (ONT).

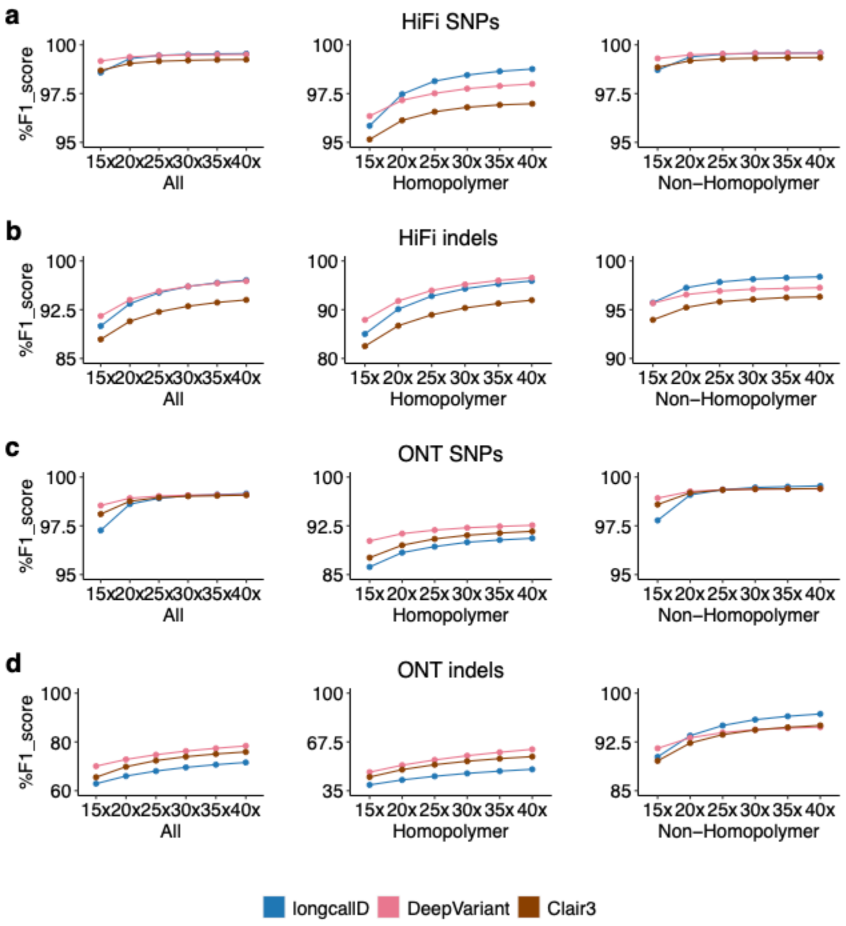


**Supplementary Fig. 2 | Small variant calling F1 scores across different read coverages.** a, F1 scores for SNPs (HiFi and T2T). b, F1 scores for indels (HiFi and T2T). c, F1 scores for SNPs (ONT). d, F1 scores for indels (ONT).

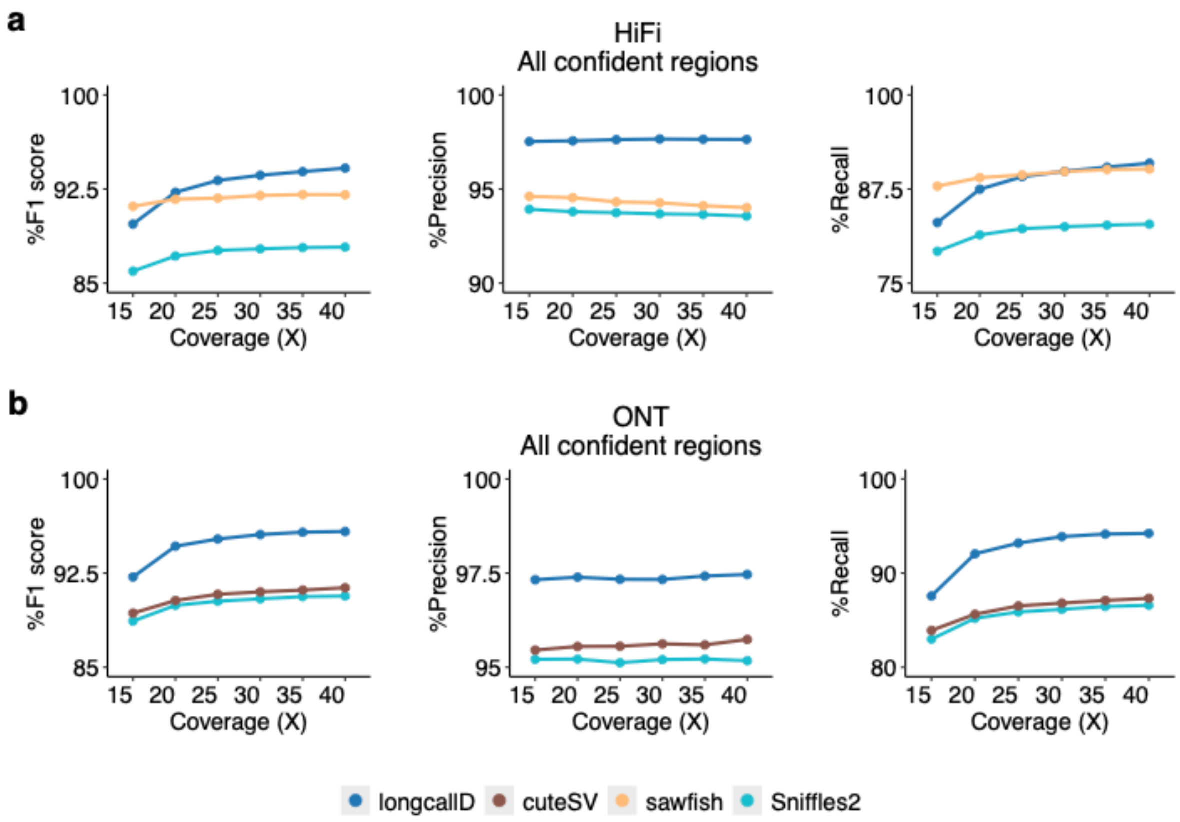


**Supplementary Fig. 3 | SV calling F1 scores across different read coverages.** a, F1 scores on HiFi data. b, F1 scores on ONT data.

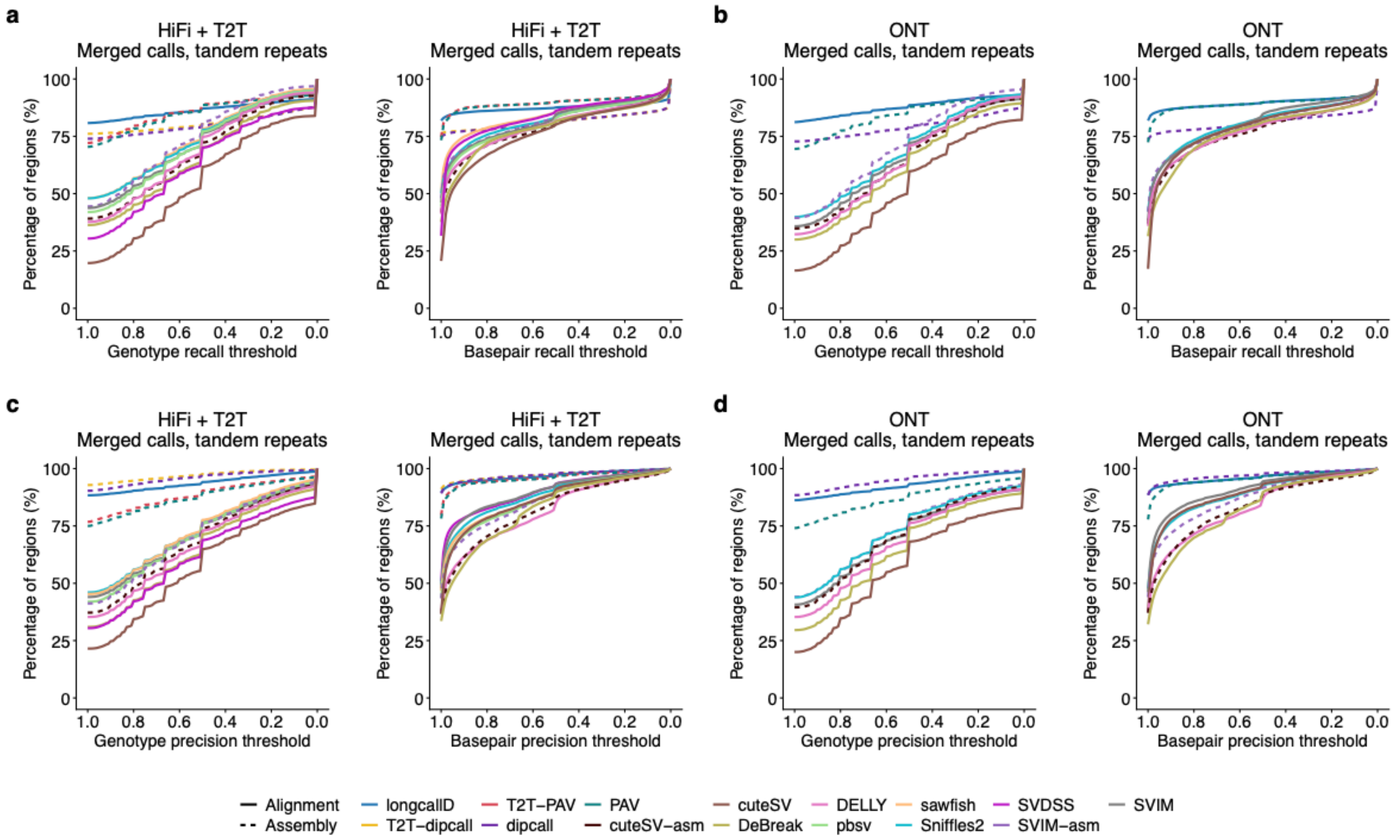


**Supplementary Fig. 4 | Region-wise genotype and basepair recalls and precisions.** a, Region-wise recall distribution in tandem repeats (HiFi and T2T), merged calls are shown for SV-only callers. b, Region-wise recall distribution in tandem repeats (ONT). c, Region-wise precision distribution in tandem repeats (HiFi and T2T). d, Region-wise precision distribution in tandem repeats (ONT).

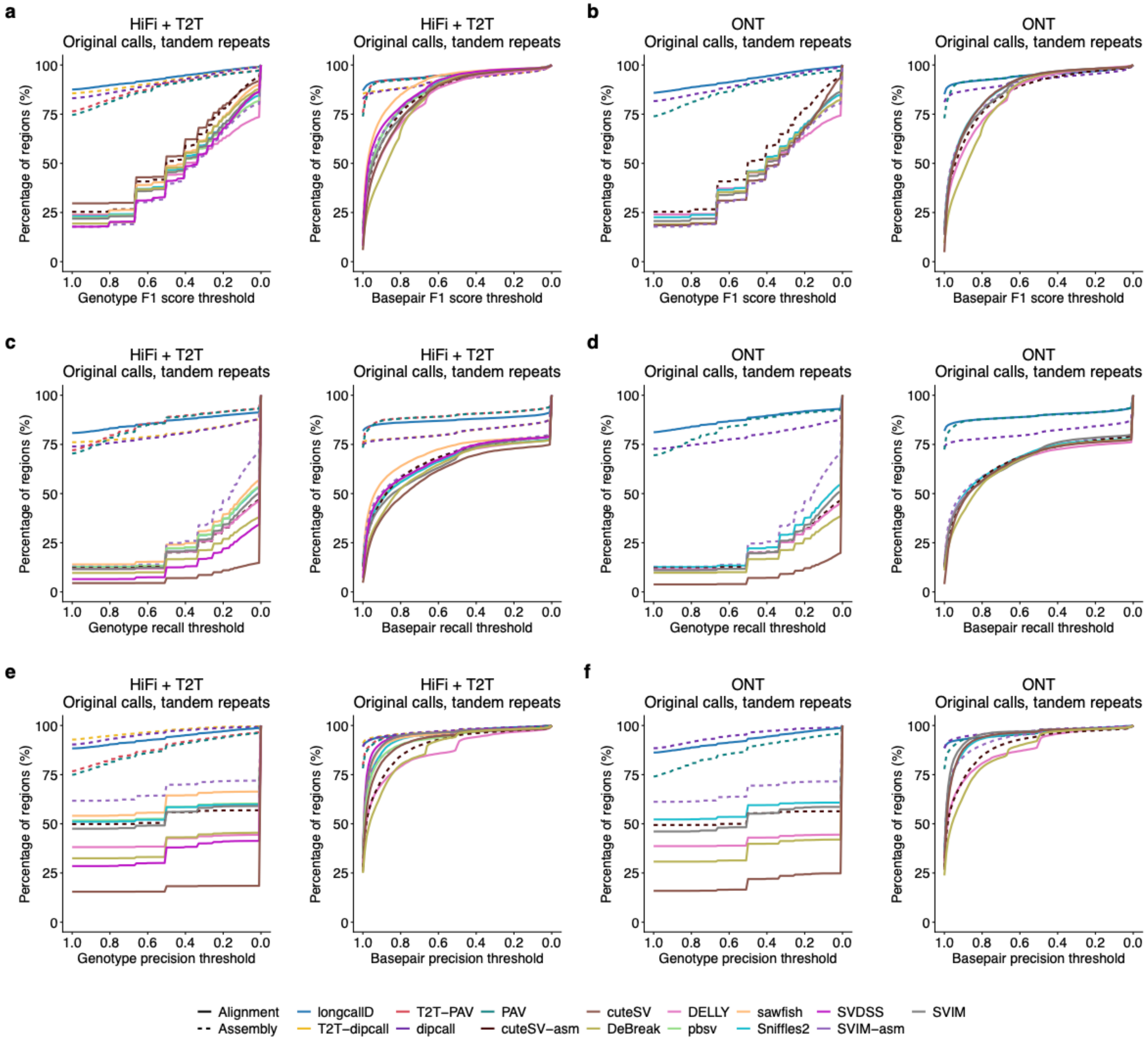


**Supplementary Fig. 5 | Region-wise genotype and basepair F1 scores, recalls and precisions.** a, Region-wise F1 score distribution in tandem repeats (HiFi and T2T), original calls are shown for SV-only callers. b, Region-wise F1 score distribution in tandem repeats (ONT). c, Region-wise recall distribution in tandem repeats (HiFi and T2T). d, Region-wise recall distribution in tandem repeats (ONT). e, Region-wise precision distribution in tandem repeats (HiFi and T2T). f, Region-wise precision distribution in tandem repeats (ONT)

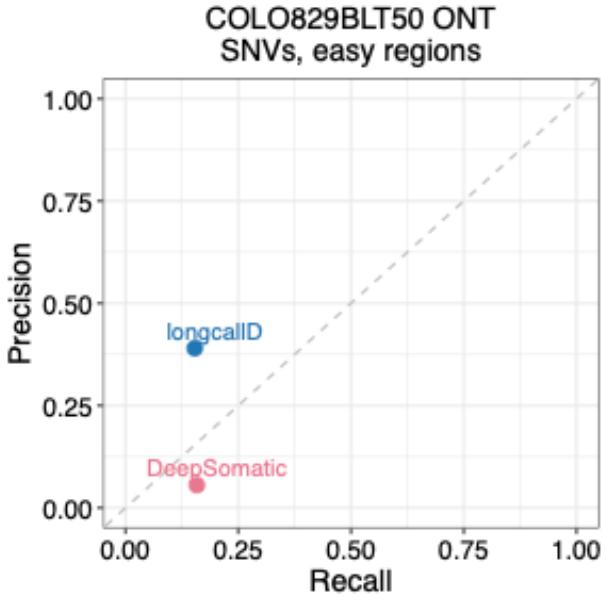


**Supplementary Fig. 6 | Precision and recall of mosaic SNV calls on COLO829BLT50 in easy regions using ONT data.** Clair-Mosaic was excluded as no SNV calls were output.

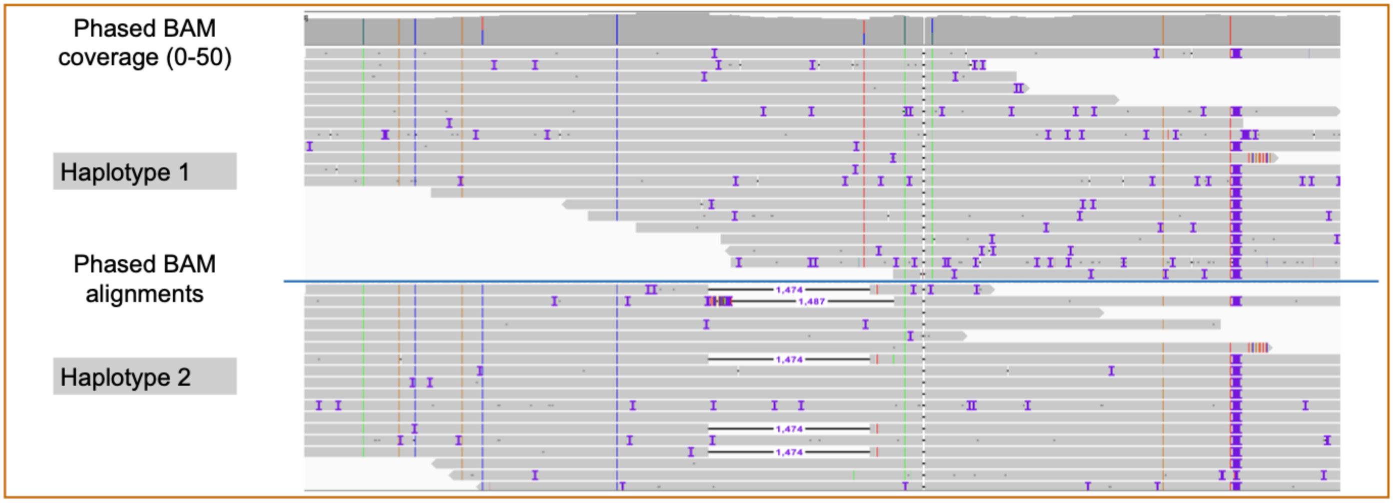


**Supplementary Fig. 7 | Fully validated mosaic deletion identified from HG002 HiFi data, detected by both longcallD and Sniffles2.** Reads were aligned to GRCh38 reference. A 1474 bp deletion is supported by five long reads, all originating from haplotype 2.
